# Inhibitors of ROCK kinases induce multiple mitotic defects and synthetic lethality in BRCA2-deficient cells

**DOI:** 10.1101/2022.06.24.497514

**Authors:** Julieta Martino, Sebastián O. Siri, Natalia S. Paviolo, Cintia Garro, María F. Pansa, Sofía Carbajosa, Aaron C. Brown, José L. Bocco, Israel Gloger, Gerard Drewes, Kevin P. Madauss, Gastón Soria, Vanesa Gottifredi

## Abstract

BRCA2-deficient cells are highly sensitive to poly-ADP-ribose polymerase inhibitors (PARPi) due to their impaired homologous recombination repair. This increased cytotoxicity is triggered by DNA replication stress induced by PARP trapping on DNA. Thus, it is broadly assumed that DNA damage is a prerequisite for BRCA2 synthetic lethality (SL). Here we show that inhibiting ROCK kinases in BRCA2 deficient cells, triggers SL independently from acute replication stress. In contrast, such SL is preceded by enhanced M-phase defects such as anaphase bridges, and abnormal mitotic figures, which were associated with multipolar spindles, supernumerary centrosomes and multinucleation. SL was also triggered by inhibiting Citron Rho-interacting kinase, another enzyme which, similarly to ROCK kinases, regulates cytokinesis. Together, these observations suggest cytokinesis failure as trigger of mitotic abnormalities and SL in BRCA2 cells. Furthermore, preventing mitotic entry by Early mitotic inhibitor 1 (EMI1) depletion promoted survival of BRCA2 deficient cells treated with inhibitors of ROCK kinases, thus reinforcing the association between M-phase and the cell death in BRCA2 deficient cells. This novel mechanism of SL induction is in contrast to the one triggered by PARPi and uncovers mitosis as an Achilles heel of BRCA2 deficient cells.

## Introduction

Hereditary breast and ovarian cancer (HBOC) is an autosomal dominant disease that accounts for 5-10% of breast (Krainer et al., 1997; Langston, Malone, Thompson, Daling, & Ostrander, 1996) and 15% of ovarian cancer cases (Pal et al., 2005; Zhang et al., 2011). HBOC is primarily caused by mutations in the breast cancer susceptibility genes BRCA1 and BRCA2 (Futreal et al., 1994; Miki et al., 1994; Wooster et al., 1995). BRCA1 and BRCA2 are DNA repair genes and their protein products regulate homologous recombination (HR), a repair pathway that is recruited to highly toxic DNA double-strand breaks (DSBs) (Prakash, Zhang, Feng, & Jasin, 2015). BRCA1 and BRCA2-deficient cells exhibit structural chromosome abnormalities and are highly sensitive to DNA damaging agents (Moynahan, Cui, & Jasin, 2001; Patel et al., 1998; Yu et al., 2000). Additionally, BRCA-deficient cells exhibit translocations, large deletions and chromosome fusions (Moynahan et al., 2001; Yu et al., 2000). This chromosome instability underlies the tumorigenicity of BRCA-deficient tumors and underscores the important tumor suppressor function of BRCA genes in cells.

Mutations in BRCA genes are highly penetrant and their carriers have high risk of developing early onset breast and ovarian cancer (Antoniou et al., 2003; King, Marks, & Mandell, 2003). Carriers of BRCA mutations are also at an increased risk of developing other tumor types such as pancreas, prostate and melanoma (Cavanagh & Rogers, 2015; Gumaste et al., 2015). BRCA-deficient patients whose mutations are detected before cancer onset are suggested to undergo highly invasive surgeries such as salpingo-oophorectomy and mastectomy. The standard of care for BRCA patients with tumors is similar to the approach used for patients with sporadic tumors, with the exception of some types of BRCA tumors which might be more sensitive to platinum-based therapies (Vencken et al., 2011; Yang et al., 2011). Unfortunately, chemotherapy resistance to platinum agents is common and alternative therapies are most needed for these patients.

One alternative therapy, already available clinically, are poly-ADP-ribose polymerase (PARP) inhibitors which are highly effective in killing BRCA-deficient cells (Bryant et al., 2005; Farmer et al., 2005; McCabe et al., 2006) and several PARP inhibitors (PARPi) have been approved for clinical use. The synthetic lethality (SL) observed between BRCA deficiency and PARPi is due to the ability of PARPi to physically trap PARP on the DNA (Murai et al., 2014; Murai et al., 2012). PARP trapping triggers DNA replication stalling and collapse, which require HR-mediated repair, a mechanism that is facilitated by BRCA1 and BRCA2 and which is therefore impaired in BRCA1 and BRCA2-deficient cells (Prakash et al., 2015). As with every cancer therapy, resistance to PARPi is also observed in the clinic (Barber et al., 2013). Molecular mechanisms of resistance to PARPi include but are not limited to secondary mutations that restore HR function, increased drug efflux, and decreased PARP trapping (D’Andrea, 2018; Noordermeer & van Attikum, 2019).

Although BRCA proteins were originally described for their key roles within HR, we currently know that BRCA1 and BRCA2 have pleiotropic functions, performing other functions outside canonical HR (Lee, 2014; Venkitaraman, 2014). Thus, it is likely that multiple targets not restricted to HR could be exploited for SL therapeutic approaches. This concept has been corroborated for BRCA1 in a phenotypic screening in which we tested BRCA-deficient cells for SL against the kinase inhibitor library PKIS2 (Carbajosa et al., 2019). Our findings unveiled that BRCA1 cells are highly sensitive to inhibition of Polo-like kinase 1 (PLK1) and that this sensitivity does not require excess DNA damage caused by external agents.

In this study, we present findings indicating that BRCA2-deficient cells are highly sensitive to the inhibition or depletion of ROCK kinases (ROCK), which regulate actin cytoskeleton dynamics. Unlike PARPi, ROCK inhibitors (ROCKi) did not induce acute replication stress in BRCA2-deficient cells but instead triggered mitosis defects including cytokinesis failure, polyploidy, aberrant multipolar spindles and centrosome amplification. Remarkably, SL-induction was also observed after inhibition of Citron Rho-interacting kinase (CITK), an enzyme that regulates cytokinesis at the level of mitotic furrow cleavage, indicating that cytokinesis failure is likely the trigger of this novel SL interaction. Moreover, preventing mitotic entry via depletion of Early mitotic inhibitor 1 (EMI1), abrogated ROCKi-induced cell killing. In conclusion, while the accumulation of DNA damage in S phase is required for PARPi-mediated cell death (Chaudhuri et al., 2016; Schoonen et al., 2017), our findings highlight that BRCA2-deficient cells bear additional vulnerabilities outside S phase that could represent promising new SL targets.

## Results

### BRCA2–deficient cells are sensitive to ROCK inhibition

In a previous work (Carbajosa et al., 2019) we developed a phenotypic survival screening method to evaluate the differential sensitivity of BRCA1-deficient cells against 680 ATP-competitive kinase inhibitors provided by GlaxoSmithKline (Drewry et al., 2017; Elkins et al., 2016). Briefly, the screening was performed using HCT116^p21-/-^ cell lines in which BRCA1 or BRCA2 were stably downregulated using shRNA (Figure 1A). This allowed comparison of BRCA-proficient vs. BRCA-deficient cell lines on an isogenic background that is easy to grow and tolerates seeding at densities that allow long term (i.e.: 6 days) survival analysis. Additionally, we used a p21 knockout background, which attenuates the cell cycle arrest that otherwise would mask the cytotoxic phenotypes during the screening time frame.

**Figure 1.**
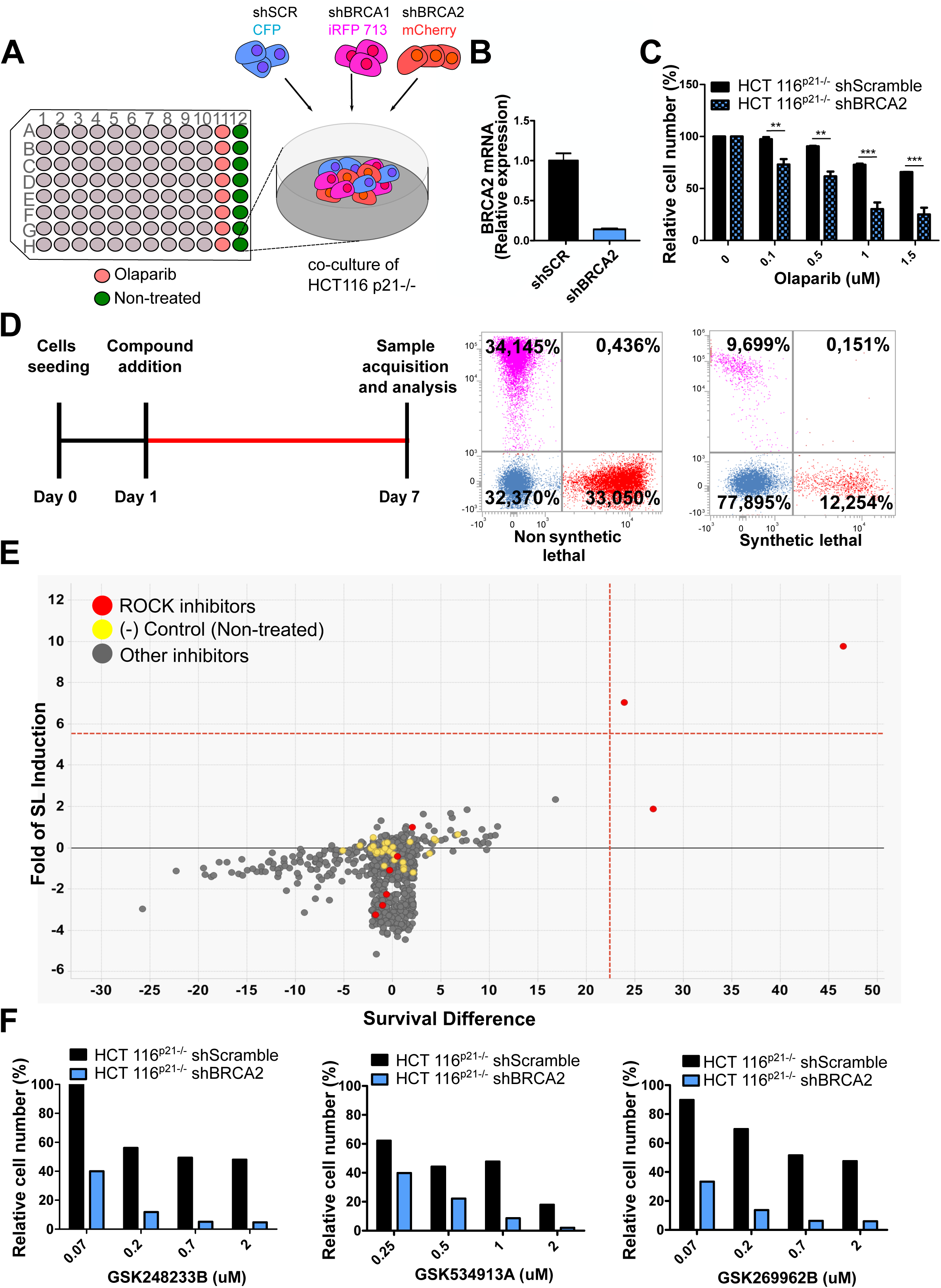
Phenotypic screening identifies ROCK kinases as potential targets for synthetic lethality in BRCA2 cells. **A)** The screening assay consists in the co-culture of isogeneic BRCA-proficient and BRCA-deficient cell lines in equal proportions on each well of 96-well plates. For this, double stable cell lines tagged with different fluorescent proteins (CFP, iRFP, and mCherry) and expressing shRNAs for Scramble, BRCA1 or BRCA2 were generated as described in (Carbajosa et al., 2019). **B)** Quantitative real-time PCR of BRCA2 in shScramble and shBRCA2 HCT116^p21-/-^ cells (N=2). **C)** Relative cell number (%) of HCT116^p21-/-^ cells expressing shScramble and shBRCA2 and treated with the indicated concentrations of Olaparib (N=2). D) Schematic timeline of the screening protocol and examples of results from each well. A tested compound can be “non synthetic lethal” (the ratio between the populations’ percentage remains unchanged when compared to the ratio used for seeding-∼33% for each cell line); or “synthetic lethal” (the ratio between cell types is altered when compared to the ratio used for seeding, with selective depletion of cells within the BRCA1- and/or BRCA2-deficient populations). E) Screening results of PKIS2 library compounds (0.1 uM) in shBRCA2 HCT116^p21-/-^ cells. Compounds were plotted based on their fold of SL (y axis) and their survival difference (x axis). A compound was considered a “hit” if it exhibited a >5 standard deviation on these two variables. Fold of SL (y axis): the ratios of the different populations in each individual well. Survival difference (x axis): compares treated cells with the untreated control in the same plate. ROCK inhibitors and other inhibitors are plotted in red and grey, respectively. For statistical analysis of the screening, please refer to Carbajosa et al. (2019). F) Relative cell number (%) of shScramble and shBRCA2 HCT116^p21-/-^ cells at different ROCK inhibitors. Statistical analysis was performed with a two-way ANOVA followed by a Bonferroni post-test (*p<0.05, **p<0.01, ***p<0.001). Data in B-C are shown as the average of 2-3 independent experiments with the standard error of the mean.

In this work, we analyzed the screening results of the BRCA2-deficient cell population. BRCA2 depletion by shRNA in HCT116^p21-/-^ cells was sufficient to trigger increased sensitivity to Olaparib (Figure 1B-C). For the analysis, we focused on compounds that induced SL exclusively in the BRCA2-deficient population and were not toxic to control samples or BRCA1-deficient cells (Figure 1D). Interestingly, BRCA2 deficient cells showed remarkable sensitivity to three inhibitors of ROCK kinases (ROCK) (Figure 1E and Figure 1-figure supplement 1A). The selective activity of each compound was further validated in a dose-response curve (Figure 1F). To test the sensitivity of BRCA2-depleted cells to ROCK inhibition, we took advantage of two commercially available ROCK inhibitors (ROCKi), Fasudil and Ripasudil, which are approved for diseases other than cancer (Garnock-Jones, 2014; Shi & Wei, 2013). Both are ATP-competitive inhibitors targeting ROCK1 and ROCK2 (Nakagawa et al., 1996). We performed survival assays with Fasudil in several cellular models of BRCA2 deficiency including the HCT116^p21-/-^ cell line used in the screening (Figure 2A). We also tested survival in DLD-1/DLD-1^BRCA2-/-^ paired cell lines, which are BRCA2 knockout (Figure 2B) and the PEO4/PEO1, V-C8 #13/V-C8 paired cell lines (see description of cell lines in the methods section - Figure 2C-D). SL was observed in all BRCA2-deficent cell line models following Fasudil treatment (Figure 2A-D). Cell death was confirmed using SYTOX green, a dye that only enters cells when cellular membranes have been compromised (Figure 2E). Similar differences between control and BRCA2-deficient counterparts were observed with Ripasudil, another ROCKi (Figure 1-figure supplement 1B-C). In contrast, the BRCA1-deficient cell line HCC1937 (Tomlinson et al., 1998), which is sensitive to Olaparib (Figure 1- figure supplement 1D), did not exhibit increased sensitivity to Fasudil or Ripasudil compared to the complemented HCC1937^BRCA1^ cell line (Treszezamsky et al., 2007) (Figure 1- figure supplement 1E-F). Similar results were observed using HCT116 cellular models depleted from BRCA1 (Figure 1- figure supplement 1G-I). The unique sensitivity of BRCA2-deficient cells to ROCKi suggests that the SL observed is likely independent of the homologous recombination function of BRCA2.

**Figure 2.**
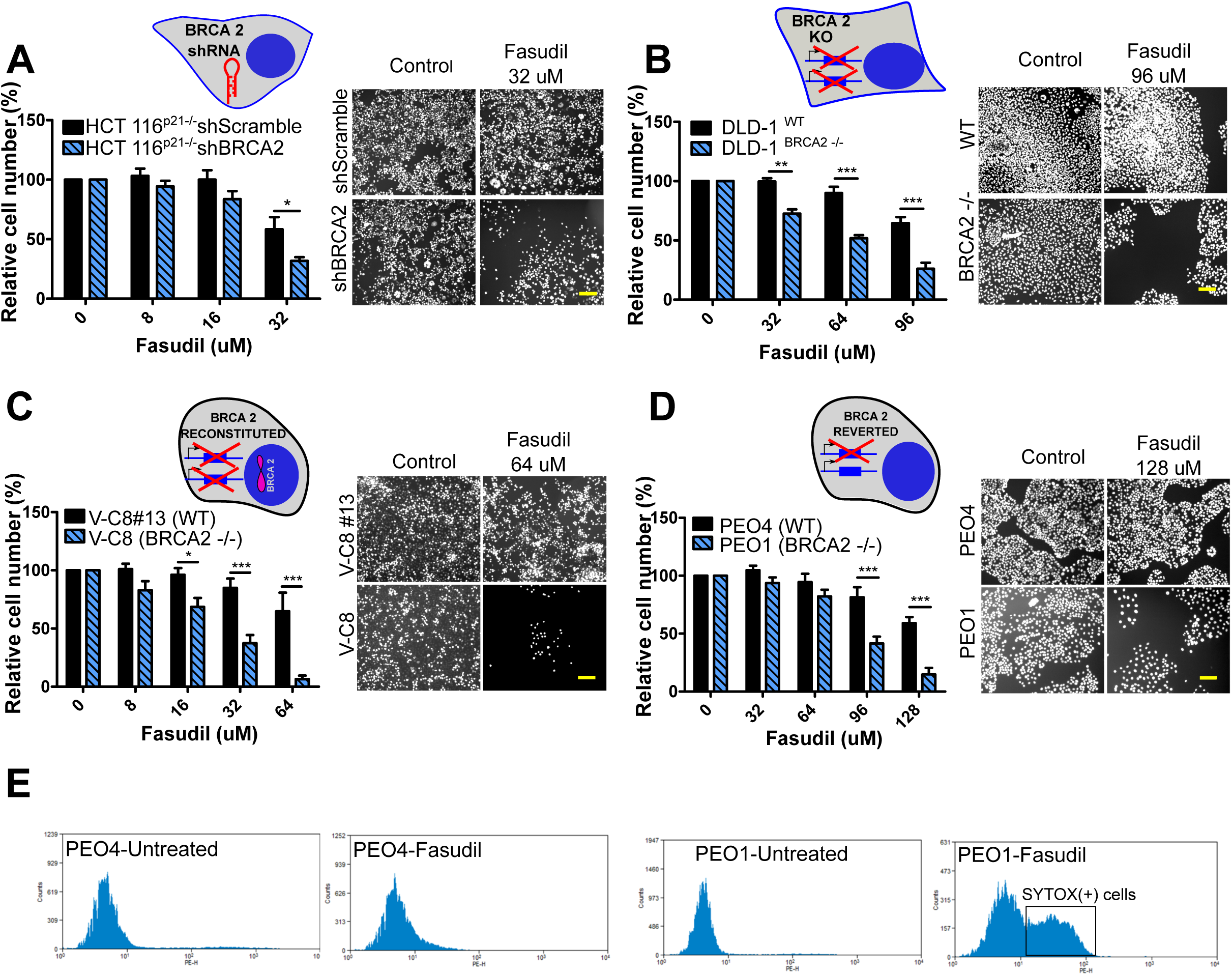
BRCA2-deficient cells are selectively killed by the ROCK kinase inhibitor Fasudil. **A)** Relative cell number (%) of shScramble and shBRCA2 HCT116^p21-/-^ cells after 6 days of treatment with Fasudil (N=3). **B)** Relative cell number (%) of DLD-1^WT^and DLD-1^BRCA2-/-^ after 6 days of treatment with Fasudil (N=2). **C)** Relative cell number (%) of V-C8#13 and V-C8 cells after 6 days of treatment with Fasudil (N=3). **D)** Relative cell number (%) of PEO4 and PEO1 cells after 6 days of treatment with Fasudil (N=2). Panels A-D: the cell cartoon shows the BRCA2 status of the modification introduced at last in each pair of cell lines (see Material and Methods for further details). Black borders indicate that the modification generated a BRCA2 proficient status and blue borders indicates BRCA2 deficiency. **E)** FACS analysis of SYTOX green-stained PEO4 and PEO1 cells 6 days after Fasudil treatment (128 uM, N=2). Statistical analysis was performed with a two-way ANOVA followed by a Bonferroni post-test (*p<0.05, **p<0.01, ***p<0.001). Data in A-D are shown as the average of 2-3 independent experiments with the standard error of the mean.

Importantly, we observe strong SL by ROCKi in growing conditions that triggered only mild sensitivity to PARPi. While HCT116^p21-/-^ shBRCA2, V-C8 and DLD-1^BRCA2-/-^ were all sensitive to Olaparib (Figure 2- figure supplement 1A), PEO1 showed only modest sensitivity to Olaparib in our experimental conditions (Figure 2- figure supplement 1B), despite reports indicating that they should depict sensitivity to PARPi (Sakai et al., 2009; Stukova et al., 2015; Whicker, Lin, Hanna, Sartorelli, & Ratner, 2016). We confirmed that PEO1 did not express BRCA2. The BRCA2 mutation in PEO1 (5193C>G) creates a premature stop codon and also a digestion site for the enzyme DrdI. In contrast, the reversion mutation in PEO4 (5193C>T) abolishes this site (Figure 2- figure supplement 1C). Consistent with their expected point mutation, following DrdI digestion PEO1 cells showed two DNA fragments (480 bp and 214 bp), which were not observed in PEO4 cell lines (Figure 2- figure supplement 1D). Additionally, as previously reported for BRCA2 deficient cell lines, (Sakai et al., 2009; Stronach et al., 2011; Stukova et al., 2015; Whicker et al., 2016) PEO1 cells are sensitive to cisplatin (Figure 2- figure supplement 1E). Our results suggest that while clonogenic assays and other approaches may better expose the sensitivity of PEO1 to Olaparib, strong SL induced by ROCKi is observed in growing conditions that reveal only mild sensitivity to PARPi. Hence, synthetic lethal avenues that diverge from PARPi could provide efficient therapeutic alternatives for the treatment of BRCA2-deficient cancer cells.

### Acute replication stress is not the major driver of SL between BRCA2-deficiency and ROCK inhibition

The SL observed between BRCA deficiency and PARPi takes place downstream of the accumulation of acute replication stress caused by PARP trapping on the DNA (Murai et al., 2012; Schoonen et al., 2017). As BRCA-deficient cells keep progressing across S-phase in the presence of PARPi, PARP/DNA adducts exacerbate replication stress resulting from fork stalling, gap formation and fork collapse (Kolinjivadi et al., 2017; Lemaçon et al., 2017; Mijic et al., 2017; Panzarino et al., 2021; Schlacher et al., 2011; Taglialatela et al., 2017). Consistent with those reports, the treatment of HCT116^p21-/-^ shBRCA2 cells with Olaparib caused the accumulation of replication stress markers such as 53BP1 and γ-H2A.X nuclear foci, which represent sites of DSB formation in S phase (Figure 3A-B). In strike contrast to Olaparib, no increase in 53BP1 or γ-H2A.X foci was induced by Fasudil treatment in HCT116^p21-/-^ shBRCA2 cells (Figures 3A-B). These results were also validated in PEO cells (Figure 3C). In line with the lack of replication stress, we did not observe a difference in the percent of BrdU+ cells after 3 or 6 days of Fasudil treatment (Figure 3D). Additionally, the intensity of BrdU was also unaffected (Figure 3 - figure supplement 1). These findings point toward a cell death-mechanism that is not centered on the accumulation of DNA damage in S-phase.

**Figure 3.**
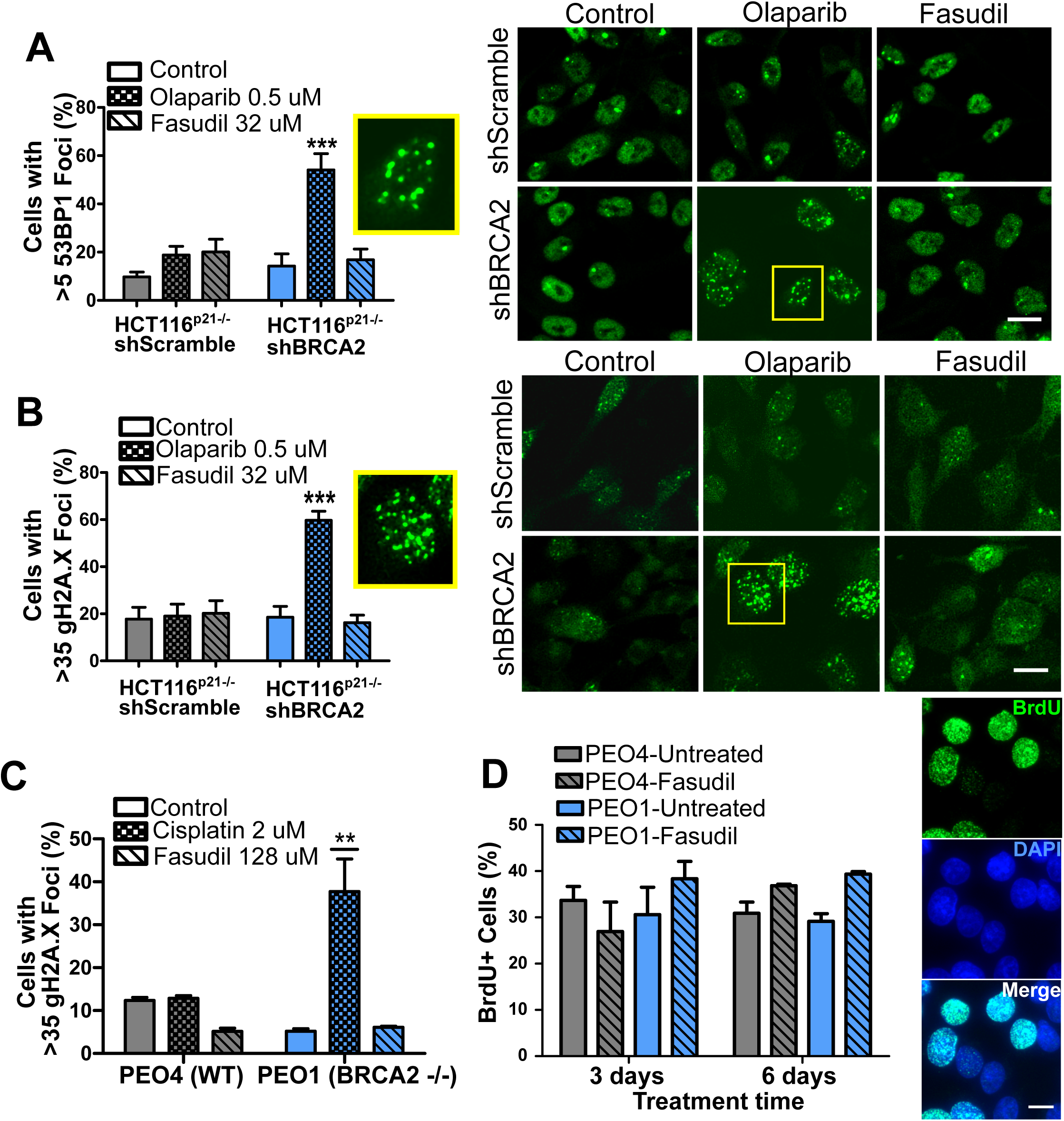
Fasudil does not induce acute replication stress in BRCA2-deficient cells. **A)** Percent of shScramble or shBRCA2 HCT116^p21-/-^ cells with >5 53BP1 foci (N=3) and representative images of the treatments. A total of 300-400 cells were analyzed per independent experiment. Cells were treated for 48 hours. **B)** Percent of shScramble or shBRCA2 HCT116^p21-/-^ cells with >35 γ-H2A.X foci (N=3) and representative images of the treatments. A total of 300-400 cells were analyzed per independent experiment. Cells were treated for 48 hours. **C)** Percent of PEO4 and PEO1 cells with >35 γ-H2A.X foci (N=2). A total of 300-400 cells were analyzed per independent experiment. **D)** Percent of PEO4 and PEO1 cells stained with BrdU after 3 and 6 days of Fasudil treatment (128 uM, N=2). A total of 500 cells were analyzed for each sample. Representative images of PEO1 cells after 3 days of Fasudil treatment (BrdU shown in green, DAPI shown in blue). Statistical analysis was performed using a two-way ANOVA followed by a Bonferroni post-test (*p<0.05, **p<0.01, ***p<0.001). Data are shown as the average of 2-3 independent experiments with the standard error of the mean.

### ROCK inhibition induces mitotic defects in BRCA2-deficient cells

To explore potential mechanisms of cell death unrelated to replication stress, we analyzed the cell cycle profiles with propidium iodide staining. Consistent with reduced survival at 6 days (Figure 2), in BRCA2-deficient cells we observed a sub-G1 peak after Fasudil treatment indicative of apoptotic cell death (Figure 4A-B). In terms of cell cycle distribution, BRCA2 cells treated with Fasudil exhibited an accumulation of cells in G2/M indicative of a G2/M arrest (Figure 4A-B). Intriguingly, BRCA2 cells also exhibited a peak of >4N polyploid cells (Figure 4A-B). By performing a detailed time course, in which samples were collected in 24-hour intervals, we observed that the polyploidy phenotype was cumulative. (Figure 4C). While the G2/M arrest in BRCA2-defcient cells appeared as early as 24 hours post-treatment, polyploidy became strongly evident at 72 hours (i.e.: 3 days). The sub-G1 population was also evident as early as 24 hours but an increase of cell death was observed at longer time points after polyploidy detection (i.e.: after 3 days). These data suggest that the accumulation of cells in G2/M precedes both polyploidy and cell death.

**Figure 4.**
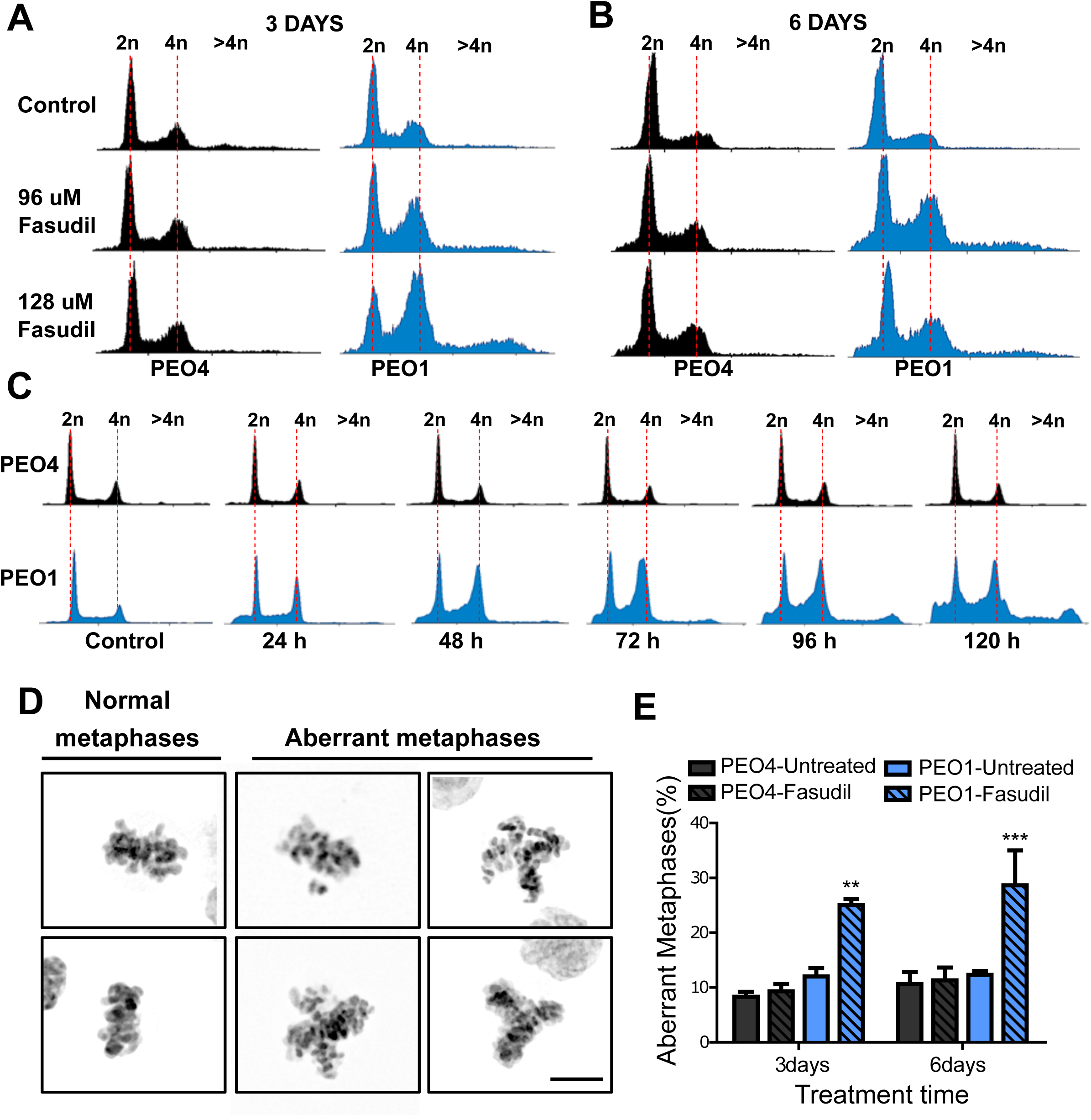
Fasudil treatment induces polyploidy and aberrant mitotic figures in BRCA2 deficient cells. **A-B)** Cell cycle analysis of PEO4 and PEO1 cells following 3 or 6 days of Fasudil treatment (96 and 128 uM) (N=3). Cells were stained with propidium iodide and DNA content was analyzed via FACS (10,000 events per sample). **C)** Cell cycle analysis of PEO4 and PEO1 cells following a time course with Fasudil treatment (N=2) (1-5 days, 64 uM). Cells were stained with propidium iodide and DNA content was analyzed via FACS (10,000 events per sample) **D)** Representative images of DAPI-stained normal and aberrant metaphases. Aberrant metaphases include metaphases with DNA being pulled in multiple directions or metaphases with misaligned chromosomes. **E)** Percent of aberrant metaphases in PEO4 and PEO1 cells 3 or 6 days after Fasudil treatment (N=3) (128 uM). A total of 100 metaphases were analyzed for each sample. Statistical analysis was performed using a two-way ANOVA followed by a Bonferroni post-test (*p<0.05, **p<0.01, ***p<0.001). Data in E are shown as the average of 2-3 independent experiments with the standard error of the mean.

The concomitant accumulation of cells in G2/M and the extra DNA content is highly suggestive of problems in the correct finalization of M phase which leads to accumulation of aberrant mitotic phenotypes. When quantifying aberrant metaphases in which the DNA was being pulled in multiple directions or in which the chromosomes were not aligned in the metaphase plate (Figure 4D) we observed a substantial increase of such phenotypes after Fasudil treatment in BRCA2-deficient cells (Figure 4E). Altogether, these data pinpoint to mitosis being dysregulated in BRCA2-deficient cells in which ROCK activity is prevented.

Aberrant metaphases can be triggered by unresolved DNA replication defects accumulated after DNA replication stress (Gelot, Magdalou, & Lopez, 2015), but can also be triggered within M phase as a consequence of aberrant mitotic spindle organization or disorganized chromosome alignment (Bakhoum, Thompson, Manning, & Compton, 2009; Shindo, Otsuki, Uchida, & Hirota, 2021; Siri, Martino, & Gottifredi, 2021). Aberrant anaphases (bridges and lagging chromosomes; Figure 5A) can also be triggered either by replication defects that are not resolved before M phase entry or intrinsic mitotic defects dissociated from S phase (Bakhoum et al., 2009; Shindo et al., 2021). We documented an increase in chromosome bridges, but not in lagging chromosomes, after Fasudil treatment of BRCA2-deficient cells (Figure 5B-C). To validate the increment of chromosome bridges observed with Fasudil, we used commercially available siRNAs against ROCK1 and ROCK2 (Figure 5D). Similar to ROCKi, ROCK1 and ROCK2 (ROCK1/2) depletion promoted the accumulation of anaphase bridges in BRCA2-deficient cells (Figure 5E). Importantly, when resulting from unresolved replication defects, anaphase aberrations are normally accompanied by chromosome aberrations (i.e.: breaks, exchanges) and micrononuclei (Finardi, Massari, & Visintin, 2020; Utani, Kohno, Okamoto, & Shimizu, 2010). However, we did not find any indication of chromosome aberrations or micronuclei in Fasudil treated BRCA2-deficient cells (Figure 3 - figure supplement 2A-B). which suggested that the trigger for anaphase bridge formation following Fasudil treatment is a defect intrinsic to M phase.

**Figure 5.**
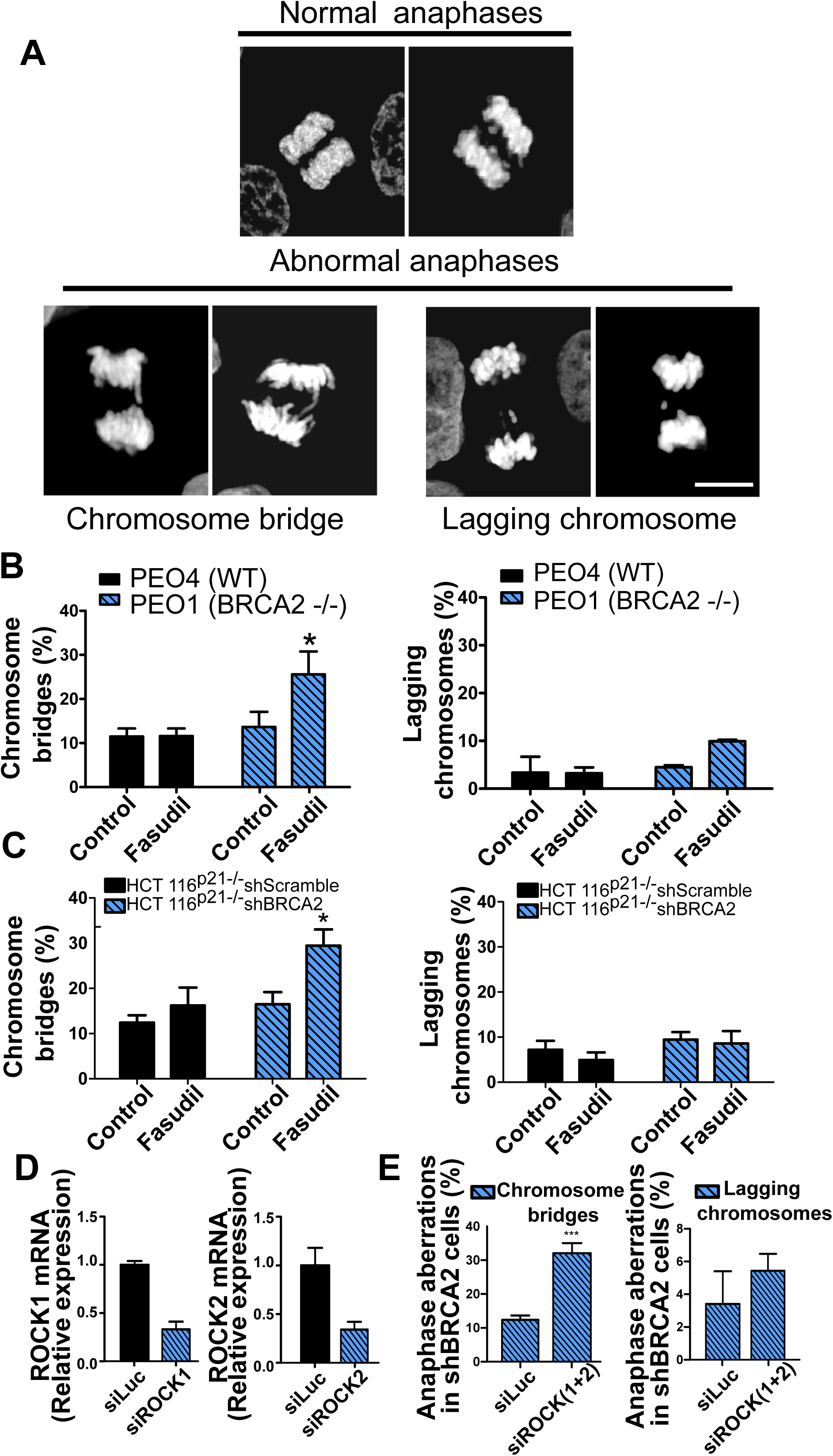
Mitotic DNA bridges accumulate in BRCA2-deficient cells following ROCK inhibition with Fasudil. **A)** Representative images of normal and abnormal anaphases with bridges and lagging chromosomes. **B)** Percentage of anaphases with chromosomes bridges and lagging chromosomes in PEO4 and PEO1 cells treated with Fasudil (128 uM). A total of 50-70 anaphases per sample were analyzed in 2 independent experiments (N=2). **C)** Percentage of anaphases with chromosomes bridges and lagging chromosome in shScramble- or shBRCA2-transduced HCT116p21-/- cells treated with Fasudil. A total of 50-70 anaphases per sample were analyzed per independent experiment (N=3). **D)** Quantitative real-time PCR of ROCK1 and ROCK2 in shBRCA2 HCT116^p21-/-^ cells transfected with 150 uM of siROCK1 or siROCK2 (N=2). E) Percentage of anaphases with chromosomes bridges and laggars in shBRCA2 HCT116p21-/- cells transfected with siROCK (1+2). A total of 50-70 anaphases per sample were analyzed per in three independent experiments (N=3). The statistical analysis of the data was performed with a two-way ANOVA followed by a Bonferroni post-test (*p<0.05, **p<0.01, ***p<0.001). Data in B-D are shown as the average of 2-3 independent experiments with the standard error of the mean.

### ROCK inhibition causes cytokinesis failure in BRCA2-deficient cells

Given that BRCA2-deficient cells treated with ROCKi accumulate M-phase defects, we explored the link between ROCK kinases and mitosis. ROCK kinases are key regulators of the actin cytoskeleton (Julian & Olson, 2014). ROCK kinases have been implicated in regulating the contraction of the actin cytoskeleton towards the end of mitosis and its downregulation or absence induces multinucleation due to cytokinesis failure (Daniels, Wang, Lee, & Venkitaraman, 2004; Jonsdottir et al., 2009; Mondal et al., 2012; Shive et al., 2010). On the other hand, BRCA2 localizes to the midbody during cytokinesis and its downregulation or absence was also reported to induce multinucleation (Lekomtsev, Guizetti, Pozniakovsky, Gerlich, & Petronczki, 2010). To explore potential roles in cytokinesis regulation between ROCK and BRCA2, we stained the actin cytoskeleton with Phalloidin to distinguish individual cytoplasms and analyzed the formation of binucleated as well as multinucleated cells after Fasudil treatment (Figure 6A). We observed a marked increase of binucleation in BRCA2-deficient cells following Fasudil treatment (Figure 6B-C). Also, we documented an increase of multinucleation in BRCA2-deficient cells transfected with siROCK (Figure 6 - figure supplement 1A-B). Consistent with the polyploidy (>4N) observed with flow cytometry, Fasudil treatment also increased the percent of multinucleated cells with 3, 4 or 5+ nuclei (Figure 6B-C). Similar to the poliploidy in the cell cycle profiles, the proportion of multinucleated cells was more severe at later endpoints (Figure 6B-C) suggesting that despite cytokinesis failure, binucleated cells continue to cycle, thus further increasing their DNA content. Indeed, the percentage of BRCA2-deficient binucleated cells transiting S phase, as revealed by cyclin A staining, was between 30-40% irrespective of ROCKi. This indicates that despite their diploid DNA content, BRCA2-deficient cells treated with Fasudil were able to start a new cell cycle and transit through a second S phase (Figure 6 - figure supplement 1C-D).

**Figure 6.**
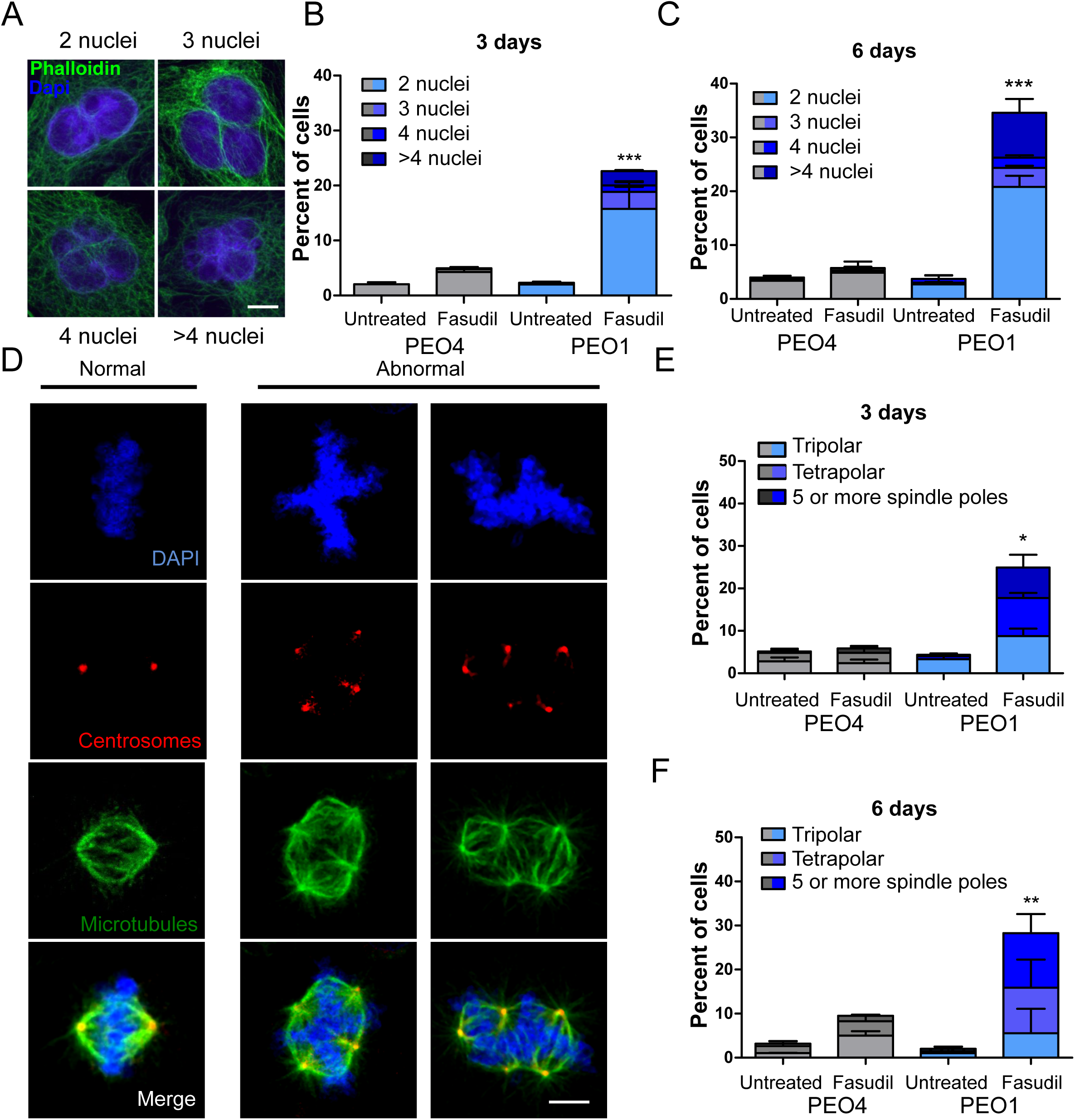
BRCA2-deficient cells exhibit cytokinesis failure, centrosome amplification and multipolar mitotic spindles following Fasudil treatment. **A)** Representative pictures of PEO1 cells after Fasudil treatment. Nuclei are stained with DAPI (shown in blue) and individual cytoplasms are stained with Phalloidin which stains the actin cytoskeleton (shown in green). **B)** Percent of binucleated and multinucleated PEO4 and PEO1 cells after 3 days of Fasudil treatment (N=3, 128 uM). **C)** Percent of binucleated and multinucleated number of PEO4 and PEO1 cells after 6 days of Fasudil treatment (N=3, 128 uM). A total of 200 cells were analyzed per sample. **D)** Representative pictures of PEO1 metaphases showing cells with normal and abnormal mitotic spindles. DNA, centrosomes, and microtubules are shown in blue, red, and green, respectively. **E)** Percent of metaphases in PEO4 and PEO1 cells with multipolar spindles after 3 days of Fasudil treatment (N=3, 128 uM). F) Percent of metaphase in PEO4 and PEO1 cells with multipolar spindles after 6 days of Fasudil treatment (N=3, 128 uM). Mitotic spindles were visualized by staining centrosomes (γ-tubulin) and microtubules (α-tubulin) and DNA was stained with DAPI. Cells were classified as having 3 spindles, 4 spindles or 5 spindles or more. A total of 100 metaphases were analyzed per sample. Statistical analysis was performed using a two-way ANOVA followed by a Bonferroni post-test (*p<0.05, **p<0.01, ***p<0.001). Data in B, C, E and F are shown as the average of 2-3 independent experiments with the standard error of the mean.

One immediate consequence of cytokinesis failure is that the resulting cell contains two centrosomes instead of one (Ganem, Storchova, & Pellman, 2007). Normal cells harbor one centrosome which duplicates only once during S phase. Duplicated centrosomes form a bipolar mitotic spindle during a normal mitosis ensuring equal chromosome distribution in daughter cells. (Nigg, 2007). In contrast, multiple centrosomes can lead to multipolar mitosis and cell death (Ganem, Godinho, & Pellman, 2009). We stained cells for gamma-tubulin and alpha-tubulin, central components of centrosomes and microtubules, respectively (Brinkley, 1997; Fuller et al., 1995) and focused on mitotic cells. BRCA2-deficient cells treated with Fasudil exhibited increased numbers of multipolar mitosis that correlated with increased centrosome number (i.e.: >2) (Figure 6D-F). Similar to previously observed phenotypes, such as aberrant metaphases, binucleated cells and polyploidy, the percent of multipolar mitosis increased at later endpoints (Figure 6F). Together, these results suggest that the cytokinesis failure and altered centrosome numbers, leads to multipolar mitosis which could be the trigger for cell death in Fasudil-treated BRCA2-deficient cells.

### Cytokinesis failure sensitize BRCA2-deficient cells to cell killing

If cytokinesis defects caused by ROCKi are the trigger of BRCA2-deficient SL, targeting other factors of cytokinesis should induce cell death as well. To test this hypothesis, we downregulated Citron Rho kinase (CITK), an enzyme that is highly enriched in the midbody during cytokinesis (Madaule et al., 1998; Sahin et al., 2019) (Figure 7A). CITK is required for proper RhoA localization at the cleavage site during late cytokinesis (Sahin et al., 2019). Similar to the phenotypes of siROCK1/2, CITK downregulation reduced cell survival of BRCA-2 deficient cells (Figure 7B and Figure 7 - figure supplement 1A). In addition, and recapitulating the effect of ROCK inhibition or depletion, CITK downregulation increased the number of multinucleated cells in BRCA2-deficient cells (Figure 7C). Most remarkably, concomitant silencing of CITK and ROCK1/2 was not additive/synergistic (Figure 7B), thus suggesting that ROCK and CITK depletion activate the same synthetic lethal mechanism in BRCA2-deficient cells. Together, these findings indicate that cytokinesis failure by multiple sources could induce death in BRCA2-deficient cells.

**Figure 7.**
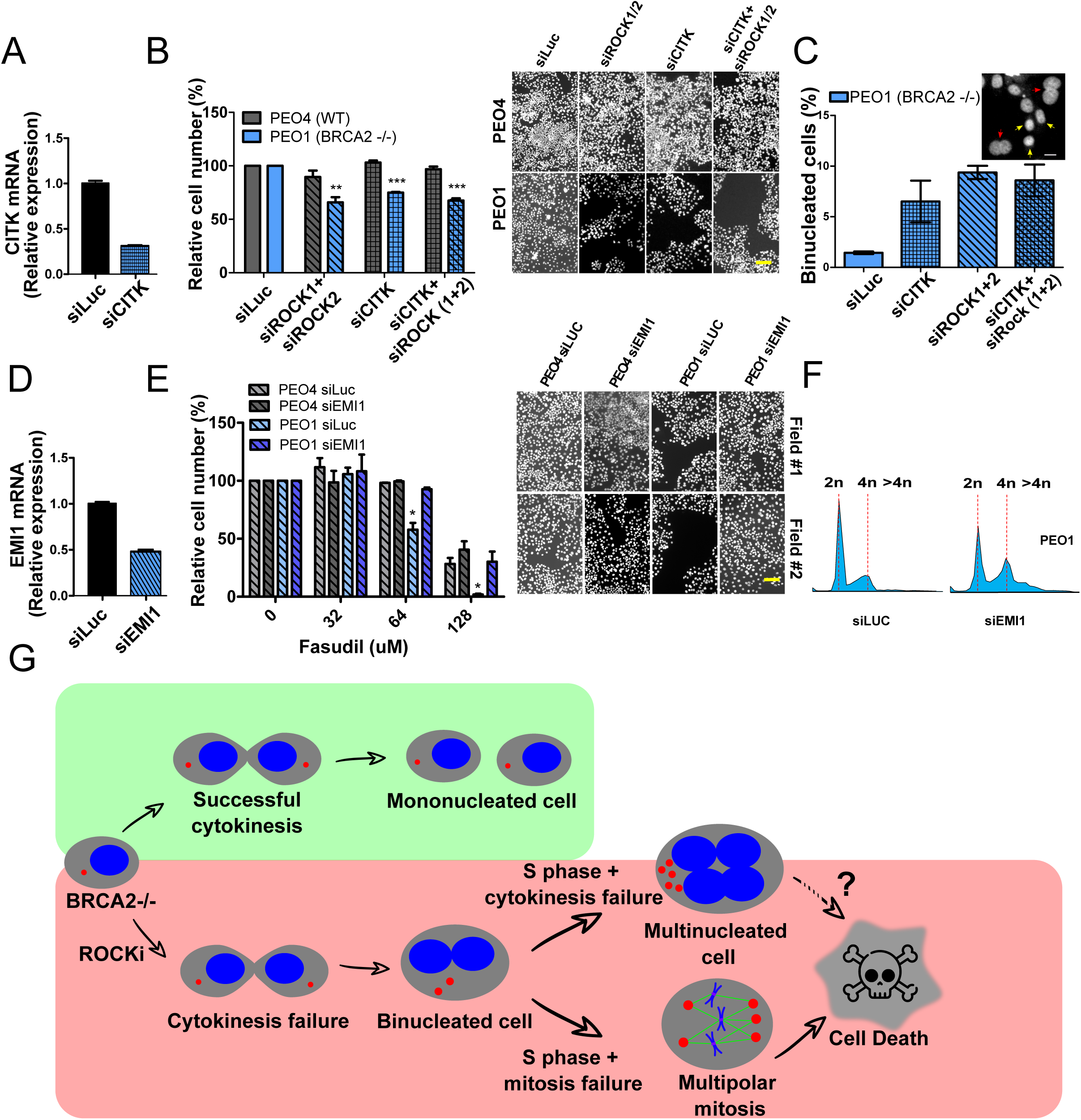
Mitosis as an alternative synthetic lethality strategy for BRCA2 deficient cells. **A)** Quantitative real-time PCR of CITK in shBRCA2 HCT116^p21-/-^ cells transfected with 150 uM of siCITK (N=2). **B)** Relative cell number (%) of PEO4 and PEO1 after 6 days of being transfected with siROCK (1+2), siCITK or siROCK (1+2)/siCITK and representative images of the transfected cells (N=2). **C)** Percent of binucleated PEO1 cells transfected with siROCK (1+2), CITK or siROCK (1+2)/siCITK (N=2). **D)** Quantitative real-time PCR of EMI1 in shBRCA2 HCT116^p21-/-^ cells transfected with 150 uM of siEMI1 (N=2). **E)** Relative cell number (%) of PEO4 and PEO1 after 6 days of being transfected with siEMI1 and treated with Fasudil (N=2). Representative images of the transfected and treated cells. **F)** Cell cycle analysis of PEO1 cells following transfection with siEMI1 for 48hs (N=2). Cells were stained with propidium iodide and DNA content was analyzed via FACS (10,000 events per sample). **G)** Model depicting the events leading to BRCA2-deficient cell death after Fasudil treatment. The inhibition or depletion of ROCK kinases in BRCA2-deficient cells leads to cytokinesis failure. As a result, the daughter cells are binucleated (4N) and have extra centrosomes (two instead of one). After DNA duplication, these cells can attempt mitosis. Mitosis entry with increased DNA content and extra centrosomes may very frequently give rise to abnormal and multipolar spindles, which leads to misaligned chromosomes and mitotic failure due to multipolar spindle formation. Alternatively, cytokinesis may fail again and cells may temporarily survive as multinucleated cells possibly facing cell death during subsequent mitotic attempts. Data in A-E are shown as the average of 2-3 independent experiments with the standard error of the mean.

If aberrant transit through mitosis is the origin of the cell death triggered by ROCKi, then the bypass of mitosis should protect those cells from cell death. To this end, we downregulated Early mitotic inhibitor-1 (EMI1), an anaphase promoting complex (APC) inhibitor that has a key role in the accumulation of mitosis activators including B-type cyclins (Reimann et al., 2001). When transfecting siEMI1, we observed a 65% reduction in EMI1 expression (Figure 7D) and, as reported by others (Robu, Zhang, & Rhodes, 2012; Shimizu et al., 2013; Verschuren, Ban, Masek, Lehman, & Jackson, 2007), accumulation of cells with G2/M DNA content or higher (Figure 7E). EMI1 depletion prevented the SL effect of ROCKi on different BRCA2 deficient cells (Figure 7F and Figure 7 - figure supplement 1B). Therefore, these results indicate that aberrant mitotic cells are likely the trigger of cell death in BRCA2-deficient cells upon ROCK inhibition.

## Discussion

### Targeting mitosis as an alternative SL strategy

In this work we used a novel screening platform developed and validated by our group (Carbajosa et al., 2019; Garcia et al., 2020) to identify ROCK as novel targets for SL induction in BRCA2 cells. Using commercially available, and clinically relevant, ROCKi (i.e.: Fasudil and Ripasudil) (Shi & Wei, 2013), we observed a dose-dependent SL-induction in multiple BRCA2-deficient cell lines which showed no signs of acute DNA replication stress. In contrast, these cells exhibited strong mitotic defects as a result of the cytokinesis failure induced by ROCKi. Remarkably, cell death by ROCK inhibition or depletion was recapitulated by the inhibition of another enzyme that facilitates cytokinesis, CITK, hence suggesting that binucleation precedes multinucleation and SL (see model in Figure 7G). In fact, robust evidence in the literature indicates that highly abnormal metaphases/anaphases, such as the ones we observed, are incompatible with cell viability (Ganem et al., 2009) and are therefore the most plausible cause the SL induced by ROCKi in BRCA2-deficient cells. While still viable, multinucleated cells are highly vulnerable. The presence of extra DNA content and centrosomes, increase the chances of abnormal spindle polarity, as well as the number of chromosomes that need to be properly aligned. In fact, attempts to trigger cell division in such states is incompatible with viability (Ganem et al., 2009; Rein, Landsverk, Micci, Patzke, & Stokke, 2015; Schoonen et al., 2017). We therefore postulate that the cytokinesis failure of a cell with 4N or more DNA content is the major driver for BRCA2 cell death following ROCK inhibition. As such, targeting mitosis alone in the absence of increased replication stress may be sufficient to kill BRCA2 cells. Future research on the mitotic functions of BRCA will certainly provide valuable information on synthetic lethal alternatives for cancers whose hallmark is the loss of this tumor suppressor gene.

### BRCA2 deficient cells can be killed in a manner that is independent from the induction of acute replication stress

In addition to the well documented replication stress-mediated toxicity of PARPi in BRCA-deficient cells (Schlacher et al., 2011; Schoonen et al., 2017), a recent report indicates that BRCA2-deficient cells can also be killed by mild replication defects which do not cause γH2A.X accumulation in S phase (Adam et al., 2021). This is dependent on the transmission of under-replicated DNA from S to M phase triggered by BRCA 1 or BRCA2 deficiency and the lack of CIP2A-TOPBP1 complex formation in M phase. In the absence of the later complex, under-replicated DNA is aberrantly processed into acentric chromosomes and micronuclei which are the source of SL (Adam et al., 2021). Our present work reveals yet another weakness of BRCA2-deficient, but not BRCA1-deficient, cells: cytokinesis. Strikingly, such SL is not preceded by the accumulation of broken chromosomes or micronuclei and is independent from canonical players of the DDR, as it is observed after ROCK or CITK inhibition.

Intriguingly, while the triggers of SL by PARPi, CIP2A-TOPBP1 complex disruption and ROCKi are remarkably different, the three mechanisms converge at mitosis ((Adam et al., 2021; Schoonen et al., 2017; Schoonen & van Vugt, 2018) and this work). CDK1 inhibition blocked micronucleation which is the trigger for BRCA-deficient cell death by CIP2A-TOPBP1 complex disruption (Adam et al., 2021), while PARPi and ROCKi-mediated cell death was abrogated by EMI1-depletion ((Schoonen et al., 2017) and this work). Hence, the transit through M phase is required for all SL events triggered in BRCA2-deficient cells. Of note, the accumulation of viable multinucleated BRCA2-depleted cells capable of enabling DNA synthesis after ROCKi reveal that, at least for a few DNA replication cycles, a cytokinesis-free cell cycle progression promotes BRCA-2 cell survival after ROCKi. Interestingly, multinucleation was reported after PARPi treatment as well (Schoonen et al., 2017) and anaphase bridges were detected both after ROCKi and PARPi as a potential source of either multinucleation or cell death ((Schoonen et al., 2017) and this work). In conclusion, despite the strong difference in the initial trigger of cell death, both after PARPi and ROCKi, BRCA2-deficient cells die at the stage of mitosis.

It should also be mentioned that our experimental analysis does not rule out that background levels of replication stress or increased levels of under-replicated DNA induced by BRCA2 deficiency could be promoting cell death by ROCK inhibition. As suggested by (Adam et al., 2021), it is possible that BRCA2-deficient cells rely more on M phase due to their defects in of the completion of DNA synthesis, making them more susceptible to suboptimal M phase (e.g.: triggered by ROCKi). However, if the source of SL was simply associated with DNA synthesis events, then it would also be likely present in BRCA1-deficient backgrounds, which we did not observe. Importantly, BRCA1 backgrounds are also vulnerable during M phase, as we previously observed SL between BRCA1 and PLK1 inhibitors (Carbajosa et al., 2019). This indicates that HR impairment is not the only possible trigger of SL in BRCA1 and BRCA2 backgrounds that could be therapeutically exploited. In the future, M phase may provide a window of opportunity for novel treatments in patients that do not respond to PARPi therapy.

### Cytokinesis failure as the trigger of the SL between BRCA2-deficiency and ROCK inhibition

We believe that DNA replication defects are not the main trigger for the SL observed with ROCKi, and that defects intrinsic to M phase are more likely account for ROCKi-induced cell death of BRCA2-deficient cells. Intriguingly, BRCA2 and ROCK functions converge at cytokinesis. ROCK kinases accumulate at the cleavage furrow (Kosako et al., 2000; Yokoyama, Goto, Izawa, Mizutani, & Inagaki, 2005), regulate furrow ingression, and their knockdown induces multinucleation (Yokoyama et al., 2005). Similarly, CITK localizes to the cleavage furrow and its downregulation or inhibition also causes multinucleation (Kosako et al., 2000; Sahin et al., 2019). Cytokinesis defects have also been reported for BRCA2-deficient cells (Daniels et al., 2004; Jonsdottir et al., 2009; Mondal et al., 2012; Rowley et al., 2011). However, BRCA2 localizes to a different cytokinesis structure than ROCK, the midbody (Daniels et al., 2004; Jonsdottir et al., 2009; Mondal et al., 2012; Rowley et al., 2011). Remarkably, previous reports suggest that the effect of BRCA2 downregulation on cytokinesis regulation may be very mild (Lekomtsev et al., 2010). Given ROCK and BRCA2 localize to cytokinesis structures that are also separated in time (furrow and midbody), the deficiency in both functions may potentiate cytokinesis failure and cell death. Supporting this, we observed not only SL but also a substantial increase in bi- or multinucleated cells when ROCK is inhibited on BRCA2-deficient backgrounds.

An alternative trigger to the SL is the formation of multipolar spindle which could be triggered by centrosome amplification. BRCA2 contributes to the regulation of centriole splitting (Saladino, Bourke, Conroy, & Morrison, 2009) and centrosome number (Ehlén et al., 2020; Saladino et al., 2009; Tutt et al., 1999). BRCA2 also localizes to centrosomes and preventing such a localization causes centrosome amplification and multinucleation (Shailani, Kaur, & Munshi, 2018). ROCK also localizes to the centrosome (Chevrier et al., 2002; Ma et al., 2006) and its activity is required for centrosome movement and positioning (Chevrier et al., 2002; Rosenblatt, Cramer, Baum, & McGee, 2004). Similar to BRCA2 deficiency, ROCK inhibition also induces centriole splitting and centrosome amplification (Aoki, Ueda, Kataoka, & Satoh, 2009; Chevrier et al., 2002; Oku et al., 2014). Interestingly, both ROCK and BRCA2 bind nucleophosmin (NPM/B23), a protein involved in the timely initiation of centrosome duplication (Ma et al., 2006; Okuda et al., 2000) and disrupting the interaction between BRCA2 and NPM/B23 induces centrosome fragmentation and multinucleation (Wang, Takenaka, Nakanishi, & Miki, 2011). Hence, the SL observed after BRCA2-deficiency and ROCKi may also be triggered by centrosome defects and lead to mitotic spindle defects, cytokinesis failure and cell death. Further work may shed additional light on this SL pathway and unravel other potential druggable targets that could be explored as therapeutic alternatives for the treatment of BRCA2-deficient tumors.

## Materials and methods

### Screening

Stable HCT116^p21-/-^ cell lines tagged with fluorescent proteins (CFP, iRFP or mCherry) and expressing Scramble, BRCA1, or BRCA2 shRNAs (Carbajosa et al., 2019) were co-cultured in equal proportions in 96-well plates for 6 days in the presence (0.1 uM) of each of the 680 compounds of the Protein Kinase Inhibitor Set 2 (PKIS2) library (Drewry et al., 2017; Elkins et al., 2016). At the end of treatment, the final cell number for each cell population was assessed with an automated flow cytometer (FACSAria II, BD Biosciences). Olaparib (#S1060, Selleckchem) at 100 nM was used a positive control in each screening plate.

For each tested compound, two scenarios are possible: A) non selective effect, where the ratio of the populations remains unchanged. The non-selective compounds can either be non-toxic (the number of cells in all populations remains the same) or toxic (the number of cells from each population decreases similarly). B) synthetic lethal: there is selective toxicity against the BRCA2 population, thus changing the relative abundance and ratio between the different populations. Additionally, a compound was considered a “hit” if it exhibited a >5 standard deviation on two values: 1) Fold of SL induction, calculated from the ratios of the different populations in each individual well; and 2) Survival difference, calculated from the differential survival when comparing a given treatment to the untreated wells in the same plate. For more extensive details on the screening platform and calculations used for the analysis please refer to Carbajosa et al. (2019).

### Lentiviral production

Lentiviral shRNA vectors were generated by cloning shBRCA2 (sequence) or shScramble (5′-GTTAACTGCGTACCTTGAGTA) into the pLKO.1-TRC vector (Grotsky et al., 2013). HEK293T cells were transfected with pLKO.1 and packaging plasmids (psPAX, and pMD2.G) 24 hours post-seeding using JetPrime transfection reagent (Polyplus). After 24 hours, media was changed. Two days after, media was collected, centrifuged and supernatants were aliquoted and stored at -80°C. Optimal viral titers were tested by serial dilutions and selected based upon minimal toxicity observed in the target cells.

### Generation of HCT116^p21-/-^ shRNA stable cell lines

HCT116^p21-/-^ cells (a kind gift from Bert Volgelstein, Johns Hopkins University) were used to generate stable shScramble or shBRCA2 HCT116^p21-/-^ cells using lentiviral transduction. For viral transduction cells were seeded in 60 mm dishes, and 24 hours post-seeding they were transduced using optimal viral titer and 8 μg/ml polybrene (#sc-134220, Santa Cruz Biotechnology). Transduced cells were selected with 1 μg/ml puromycin (#P8833, Sigma-Aldrich) 24 hours post-transduction, and grown for freezing. Frozen stocks were not used for more than three weeks after thawing. BRCA2 knockdown was confirmed using quantitative real-time PCR.

### Other cell lines and culture conditions

PEO1/PEO4: PEO1 is a BRCA2-deficient ovarian cell line derived from the ascites fluid of a patient (Langdon et al., 1988; Wolf et al., 1987). PEO4 derives from the same patient after chemotherapy resistance and has restored BRCA2 function (Sakai et al., 2009; Wolf et al., 1987). V-C8 and V-C8#13: V-C8 (a kind gift from Bernard Lopez, Gustave Roussy Cancer Center) is a BRCA2-deficient Chinese hamster lung cell line while V-C8#13 has restored BRCA2 function via one copy of human chromosome 13 harboring BRCA2 (Kraakman-van der Zwet et al., 2002). DLD-1/DLD-1^BRCA2-/-^ cell lines (# HD PAR-008 and #HD 105-00, Horizon Discovery Ltd.): DLD-1 cell lines are human colorectal cancer cell lines while the BRCA2-deficient DLD-1^BRCA2-/-^ cell line has BRCA2 exon 11 disrupted with rAAV gene editing technology (Hucl et al., 2008).

PEO4/PEO1, and DLD-1/DLD-1^BRCA2-/-^ cell lines were grown in RPMI (#31800-089, Gibco) supplemented with 10% fetal bovine serum (Natocor) and 1% penicillin/streptomycin. V-C8#13/V-C8, HCC1937^BRCA1^/HCC1937 (ATCC) and HEK293T (a kind gift from Alejandro Schinder, Fundación Instituto Leloir) were grown in DMEM (#12800082, Gibco) supplemented with 10% fetal bovine serum (Natocor) and 1% penicillin/streptomycin. All cell lines were maintained in a humidified, 5% CO_2_ incubator and passaged as needed. Cell lines were regularly checked for mycoplasma contamination. The BRCA2 and BRCA1 status of all cell lines was checked and none of the used cell lines are in the list of commonly misidentified cell lines maintained by the International Cell Line Authentication Committee.

### Drugs and treatments

Cells were treated 24 hours post-seeding. Treatment times for each experiment, ranging from 24 hours to 6 days, are specified below or in the figure legends. Olaparib (#S1060, Selleckchem) was resuspended in DMSO and stored at -20°C. ROCK inhibitors, Fasudil HCl (#A10381, Adooq) and Ripasudil (#S7995, Selleckchem) were resuspended in water and stored at -80°C. BrdU (Sigma-Aldrich) was resuspended in DMSO and stored at -20°C. BrdU-containing media (10 uM) was added to cell cultures 15 minutes before harvest. Cisplatin was resuspended in 0.9% NaCl and stored at -20°C (#P4394, Sigma-Aldrich). Cisplatin was added to cell cultures for 24 hours. All drug stocks were filter-sterilized (0.2 uM). Unless noted, all experiments were performed three times.

### Survival assay

Cells were seeded in 96 well plates and treated 24 hours post-seeding. HCT116^p21-/-^ cell lines were seeded at 1500 cells/well, V-C8 at 500 cells/well, PEO at 2500 cells/well and DLD-1/DLD-1^BRCA2-/-^ at 500 and 1500 cells/well, respectively. Each treatment had three technical replicates. Six days after treatment, plates were fixed with 2% paraformaldehyde/ 2% sucrose and stained with DAPI (#10236276001, Roche). Plates were photographed with the IN Cell Analyzer 2200 high content analyzer (GE Healthcare), using a 10x objective. A total of nine pictures per individual well were taken and all nuclei in the image were automatically counted to assess cell numbers from each well. Cell number (%) after each treatment was calculated relative to the total number of cells in untreated wells.

### Restriction enzyme digest

Genomic DNA from PEO4 and PEO1 cell lines was extracted using phenol-chloroform-isoamyl alcohol (#P3803, Sigma-Aldrich). A fragment of 694 bp within the BRCA2 gene was PCR amplified using specific primers (Forward primer: AGATCACAGCTGCCCCAAAG, Reverse primer: TTGCGTTGAGGAACTTGTGAC). PCR fragments were gel purified and equal amounts of DNA were subject to DrdI (New England Biolabs) enzyme digest following manufacturer’s instructions. Digests were run on an agarose gel and stained with ethidium bromide to visualize the band pattern.

### Chromosome aberration analysis

Cells were seeded, treated 24 hours post-treatment and 0.08 μg/ml colcemid (KaryoMAX, Invitrogen) was added 20 hours before harvest. Following trypsinization, cells pellets were incubated in hypotonic buffer (KCl 0.0075 M) at 37°C for 4 min and fixed with Carnoy’s fixative (3:1 methanol:glacial acetic acid). Cells were dropped onto slides and air-dried before staining with 6% Giemsa in Sorensen’s buffer (2:1 67 mM KH_2_PO_4_:67 mM Na_2_HPO_4_, pH 6.8) for 2 minutes. Pictures of metaphases were taken using an automated Applied Imaging Cytovision microscope (Leica Biosystems). Fifty metaphase spreads per independent experiment were analyzed for chromosome gaps, breaks and exchanges.

### Anaphase aberration assay

To quantify anaphase aberrations (bridges and lagging chromosomes), cells were fixed with 2% paraformaldehyde/ 2% sucrose for 20 min and stained with DAPI (#10236276001, Roche) to visualize anaphases. At least 50 anaphases/sample were analyzed. For image acquisition, Z-stacks were acquired with a Zeiss LSM 510 Meta confocal microscope. Maximum intensity projections were generated using FIJI (ImageJ) Imaging Software.

### Micronuclei assay

Micronuclei (MN) analysis were performed using protocols previously described previously (Federico et al., 2016). Briefly, cells were seeding at low density, treated and incubated with cytochalasin B (4.5ug/ml, Sigma-Aldrich) for 40 h. Cells were washed twice with PBS and fixed with PFA/sucrose 2% for 20 min. Phalloidin and DAPI staining were used to visualize whole cells and nuclei, respectively. 300 binucleated cells were analyzed and the frequency was calculated as MN/binucleated cells.

### Immunofluorescence

Cells were seeded on coverslips, treated, fixed for 20 min with 2% paraformaldehyde/ 2% sucrose and permeabilized for 15 min with 0.1% Triton-X 100. Following 1 hour blocking with 2.5% donkey serum in 0.05% PBS/Tween, coverslips were incubated as needed with primary antibodies: γH2A.X S139 (1:1500, #05-636-I, Millipore), 53BP1 (1:1500, #sc-22760, Santa Cruz Biotechnology), cyclin A (1:1000, #GTX-634-420, GeneTex) or Phalloidin (1:50, #A12379, Invitrogen). For BrdU staining (1:500, #RPN20AB, GE Healthcare) cells were fixed with ice-cold methanol (40 sec) and acetone (20 sec), followed by DNA denaturing in 1.5N HCl for 40 min. For staining of centrosomes (1:1000, #T6557, Sigma-Aldrich) and microtubules (1:1000, #T9026, Sigma-Aldrich) cells were fixed for 10 min with ice-cold methanol, followed by hydration with PBS. Following 1 hour of incubation with primary antibodies, cells were washed (3x/10 minutes each) with 0.05% PBS/Tween, incubated for 1 hour with anti-donkey Alexa 488 or 546 (1:200, Invitrogen), washed, stained with DAPI (#10236276001, Roche) and mounted on slides with Mowiol (Sigma-Aldrich). Slides were analyzed with 40x or 100x objectives using an Axio Observer microscope (Zeiss).

### FACS

Cells were seeded, treated and harvested at different time points (24 hours-6 days). Cells were trypsinized, fixed with ice-cold ethanol overnight, and stained with a solution of 100 μg/ml RNase (#10109142001, Roche) and 50 μg/ml propidium iodide (#P4170, Sigma-Aldrich). A total of 10,000 events were recorded using a FACSCalibur (BD Biosciences). Cell cycle distribution was analyzed with the Cytomation Summit software (Dako version 4.3). To assess cell death, cells were treated as above but following trypsinization they were stained with SYTOX Green staining following manufacturer’s instructions (#S34860, Invitrogen). A total of 10,000 events were recorded and analyzed using a FACSAria (BD Biosciences).

### Quantitative real-time PCR

Total RNA was extracted with TRIzol reagent (Invitrogen), following manufacturer’s instructions. A total of 2 ug of RNA was used as a template for cDNA synthesis using M-MLV reverse transcriptase (#28025, Invitrogen) and oligo-dT as primer. Quantitative real-time PCR was performed in a LightCycler 480 II (Roche) using the 5X HOT FIREPol EvaGreen q PCR Mix Plus (#08-24-00001, Solis BioDyne).

To calculate relative expression levels, samples were normalized to GAPDH expression. Forward (FW) and reverse (RV) primers were as follow: BRCA2 (FW: AGGGCCACTTTCAAGAGACA, RV:TAGTTGGGGTGGACCACTTG), ROCK1 (FW: GATATGGCTGGAAGAAACAGTA, RV:TCAGCTCTATACACATCTCCTT), ROCK2 (FW:AGATTATAGCACCTTGCAAAGTA, RV:TATCTTTTTCACCAACCGACTAA), CITK (FW:CAGGCAAGATTGAGAACG, RV:GCACGATTGAGACAGGGA), EMI1 (FW:TGTTCAGAAATCAGCAGCCCAG, RV:CAGGTTGCCCGTTGTAAATAGC) and GAPDH (FW:AGCCTCCCGCTTCGCTCTCT, RV GAGCGATGTGGCTCGGCTGG.

### siRNA

siRNAs were transfected using JetPrime transfection reagent (Polyplus) following the manufacturer’s instructions. Unless noted, cells were transfected for a total of 48 hours. siROCK1 (#sc-29473 Santa Cruz Biotechnology) and siROCK2 (#sc-29474, Santa Cruz Biotechnology) were used at 100 nM. siEMI1 (#sc-37611 Santa Cruz Biotechnology) and siCITK (#sc-39214 Santa Cruz Biotechnology) were both used at 100 nM.

### Statistical analysis

GraphPad Prism 5.0 was used for all statistical analyses. Regular two-way ANOVA, followed by a Bonferroni post-test or Student’s t–tests were used as appropriate. BrdU intensity was analyzed with a Kruskal-Wallis non-parametric test followed by a Dunn’s multiple comparison test. Statistical significance was set at p<0.05.

## Acknowledgements

We would like to thank Dr. Fernanda Ledda for providing critical reagents for this work. We would also like to thank all members of the Gottifredi and Soria Laboratories for insightful comments and discussions. We thank Pamela Rodriguez, Esteban Miglietta, Andrés Hugo Rossi and Carla Pascuale for technical support with tissue culture, microscopy and flow cytometry. We also thank the flow cytometry, microscopy, and cell culture facilities of CIBICI-CONICET for technical support.

## Author contributions

GS and VG conceived the original project. GS, VG and KPM supervised the study. JM, SOS, MFP, ACB, KPM, GS and VG designed the experiments. MFP and SC performed and analyzed the screening experiments. JM, SOS, CG and NP performed the rest of the experiments. All authors interpreted and analyzed the data. JM and VG designed the manuscript and flow of figures. JM and SOS generated the figures. JM and VG wrote the manuscript. All authors edited the manuscript and agreed to this description of each author’s contributions.

## Data availability

The authors confirm that the principal data supporting the findings of this study are available in Dryad-(Publish and Preserve your Data) in the next DOI https://doi.org/10.5061/dryad.bvq83bkc5. Please contact the corresponding authors for any other request.

---

**Figure 1- figure supplement 1.**
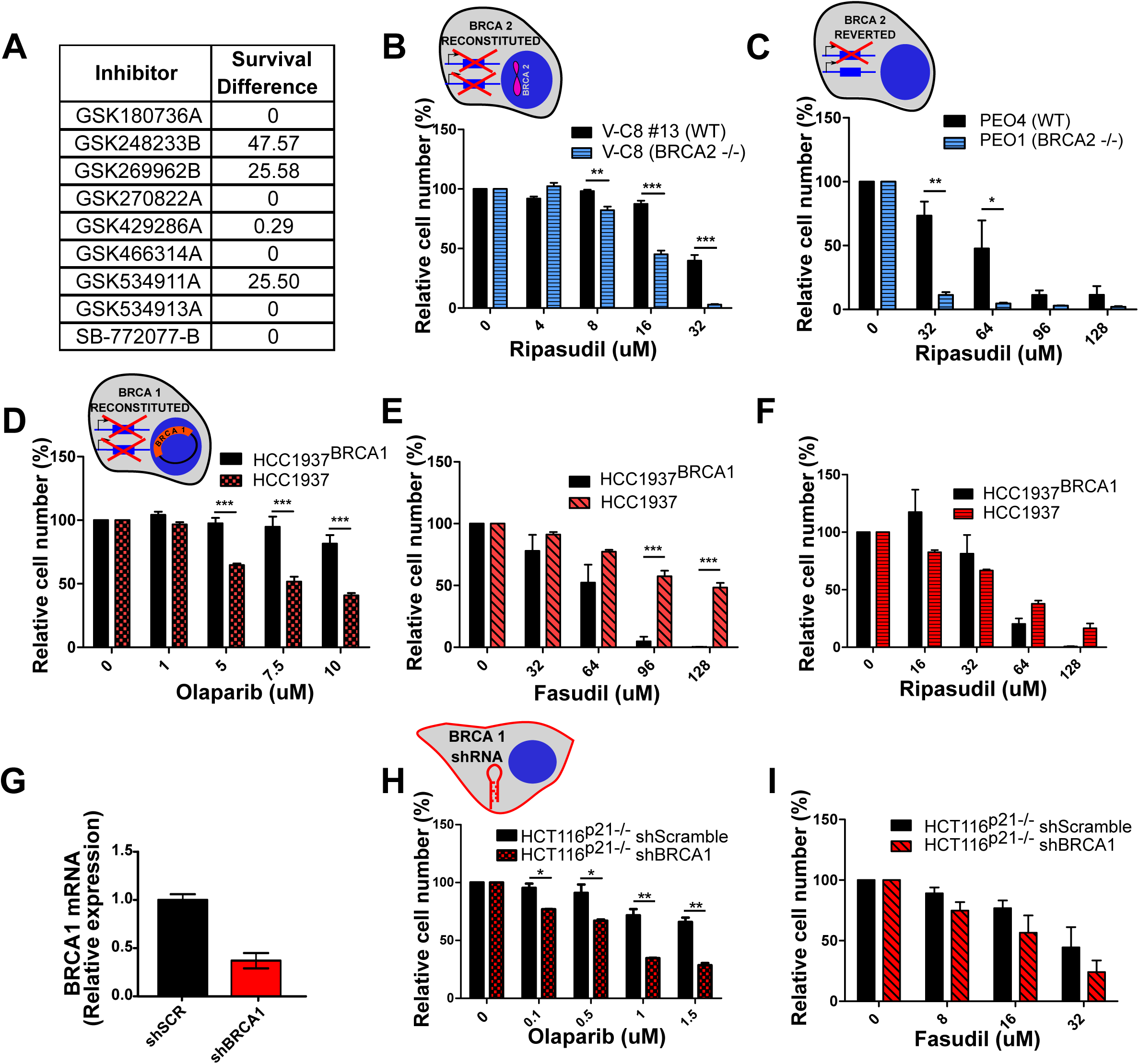
BRCA2-deficient cells are sensitive to the ROCK kinase inhibitor Ripasudil. **A)** Table listing all ROCK inhibitors from the PKIS2 library and their corresponding survival difference. **B)** Relative cell number (%) of V-C8#13 and V-C8 cells after 6 days of treatment with Ripasudil (N=2). **C)** Relative cell number (%) of PEO4 and PEO1 cells after 6 days of treatment with Ripasudil (N=2). **D)** Relative cell number (%) of HCC1937^BRCA1^ and HCC1937 cells after 6 days of treatment with Olaparib (N=2). **E)** Relative cell number (%) of HCC1937^BRCA1^ and HCC1937 cells after 6 days of treatment with Fasudil (N=3). **F)** Relative cell number (%) of HCC1937^BRCA1^ and HCC1937 cells after 6 days of treatment with Ripasudil (N=2). **G)** Quantitative real-time PCR of BRCA1 in shScramble and shBRCA1 HCT116^p21-/-^ cells. **H)** Relative cell number (%) of shScramble and shBRCA1 HCT116^p21-/-^ cells after 6 days of treatment with Olaparib (N=2). **I)** Relative cell number (%) of shScramble and shBRCA1 HCT116^p21-/-^ cells after 6 days of treatment with Fasudil (N=2). Statistical analysis was performed with a two-way ANOVA followed by a Bonferroni post-test (*p<0.05, **p<0.01, ***p<0.001). Data in B-I are shown as the average of 2-3 independent experiments with the standard error of the mean. Panel B-F and H-I: the cell cartoon shows the BRCA2 or BRCA1 status of the modification introduced at last in each pair of cell lines (see Material and Methods for further details). Black borders indicate that the modification generated a BRCA2 or BRCA1 proficient status, blue borders indicates BRCA2 deficiency and red border, BRCA1 deficiency.

**Figure 2- figure supplement 1.**
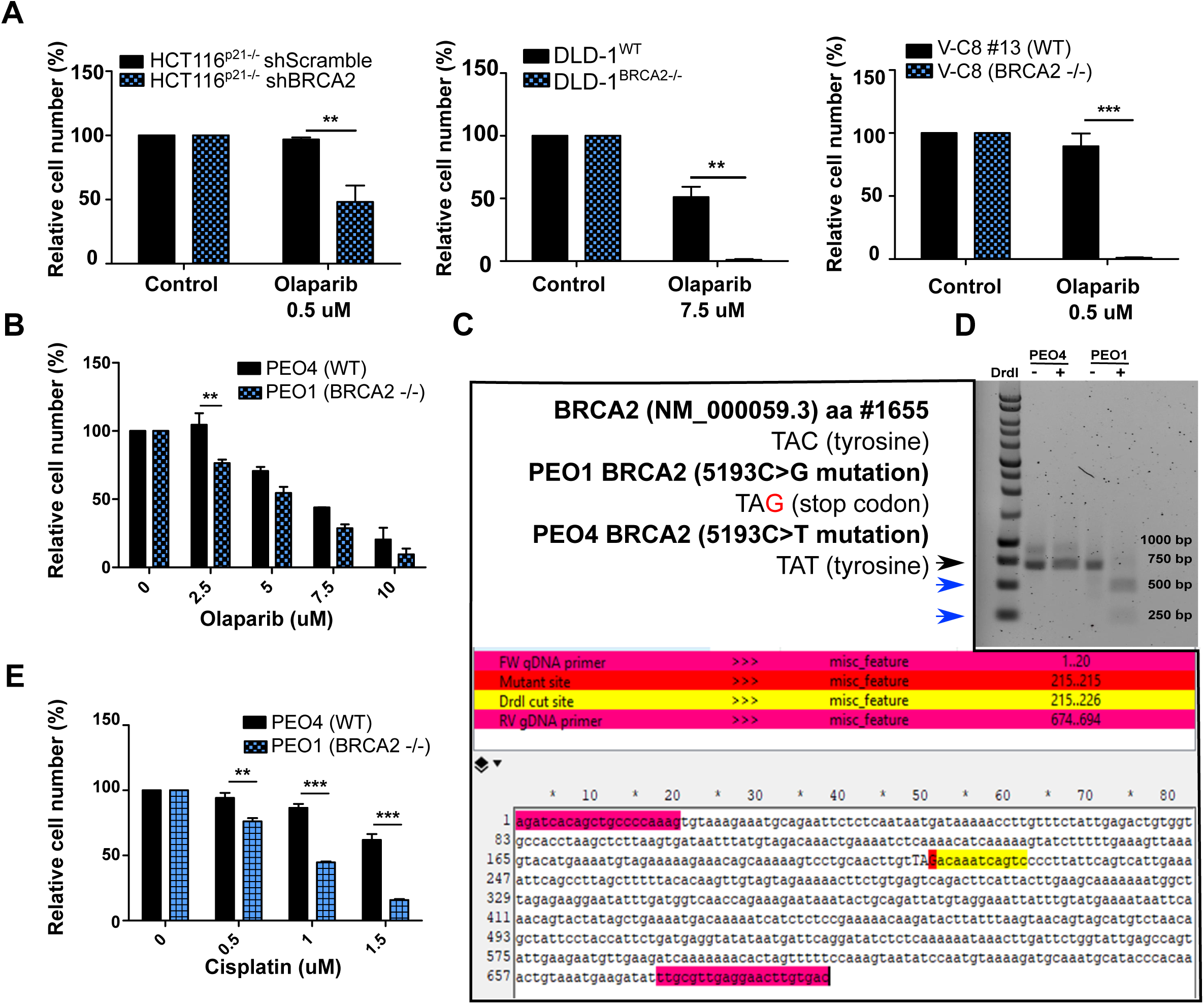
BRCA2-deficient cells are sensitive to Olaparib. **A)** Relative cell number (%) of shScramble and shBRCA2 HCT116^p21-/-^ (N=3), DLD-1^WT^ and DLD-1^BRCA2-/-^ (N=2), and V-C8#13 and V-C8 (N=4) cells after 6 days of treatment with Olaparib. **B)** Relative cell number (%) of PEO4 and PEO1 cells after 6 days of treatment with Olaparib (N=2). **C)** Nucleotide sequence (TAC, tyrosine) of aminoacid #1655 of the BRCA2 reference sequence NM_000059.3, and the nucleotide sequences found in PEO1 (TAG, stop codon) and PEO4 (TAT, tyrosine). Schematic of a fragment sequence of BRCA2 showing the primers (highlighted in magenta) used to amplify a 694 bp fragment around aminoacid #1655. The PEO1 mutation site TAG is highlighted in red. The DrdI enzyme digestion site is highlighted in yellow. **D)** Agarose gel showing undigested and digested (DrdI enzyme) products of a 694 bp fragment of the BRCA2 gene from PEO4 and PEO1 cells (Black arrow head). The nonsense mutation in PEO1 (BRCA2-deficient cell lines) generates a cut site for the DrdI enzyme giving rise to two digestion products of 480 bp and 214 bp (Blue arrow head). **E)** Relative cell number (%) of PEO4 and PEO1 cells treated with cisplatin (24 hours) followed by 5 days of growth in cisplatin-free media (N=2). Data in A, B and E are shown as the average of three technical replicates with the standard deviation. Panels A-E: Statistical analysis was performed using a t-test (*p<0.05, **p<0.01, ***p<0.001).

**Figure 3- figure supplement 1.**
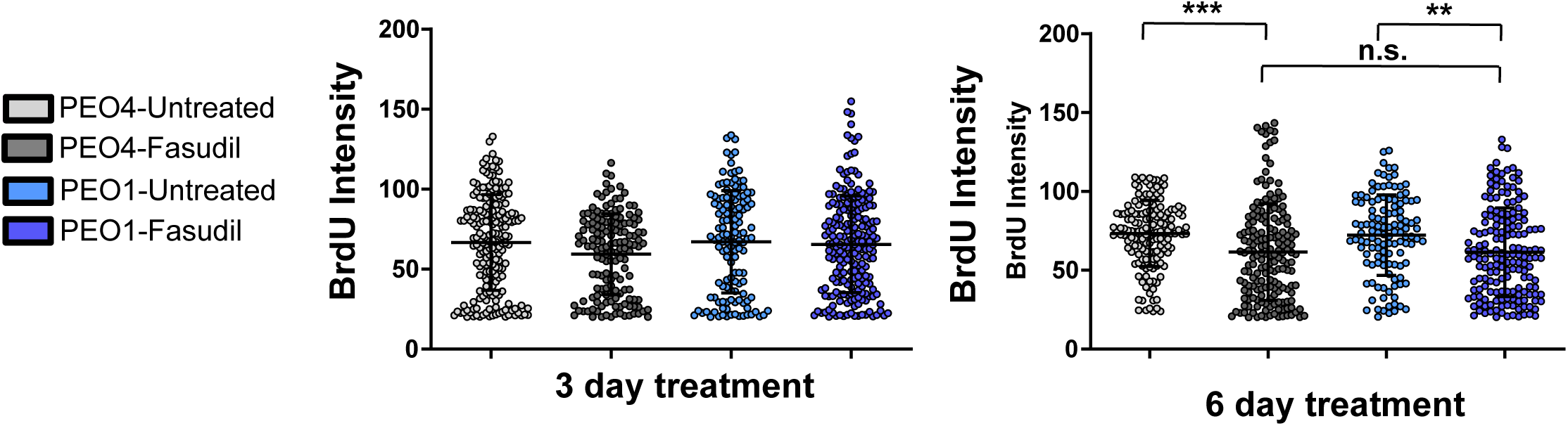
Fasudil does not alter S phase in BRCA2-deficient cells. **A)** BrdU intensity in experiments from Figure 3D. PEO4 (grey circles) and PEO1 (blue circles) cells after 3 or 6 days of Fasudil treatment (128 uM, N= 2). A total of 500 cells were analyzed for each sample. Individual intensity values per cells are displayed as a scatter plot and the average and standard deviation of each population are shown. Statistical analysis was done with a Kruskal-Wallis non-parametric test followed by a Dunn’s multiple comparison test (*p<0.05, **p<0.01, ***p<0.001).

**Figure 3 - figure supplement 2.**
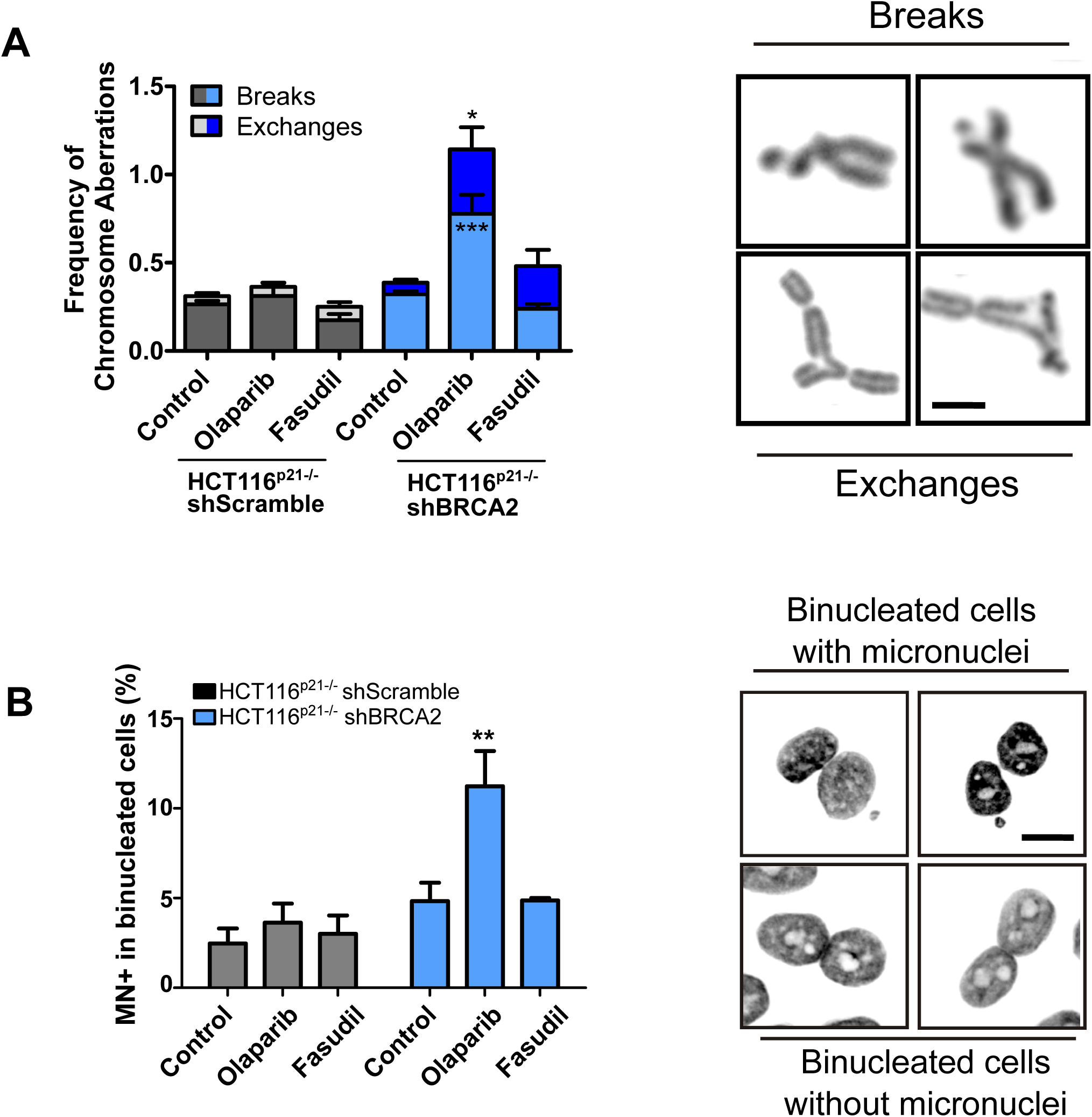
BRCA2-deficient cells treated with Fasudil do not display replication stress-derived chromosome defects. **A)** Frequency of chromosome aberrations in shScramble or shBRCA2 HCT116^p21-/-^ cells following treatment with Olaparib (0.5 uM) or Fasudil (32 uM) (N=3). A total of 50 metaphases were analyzed per condition. Chromosome aberrations include chromatid breaks and chromatid exchanges. Representative images of a chromatid break and a chromatid exchange are shown on the right. **B)** Percent of shScramble or shBRCA2 HCT116^p21-/-^ cells with micronuclei in binucleated cells (N=3). On the right, representative image of binucleated cells with and without micronuclei. 24 hours after seeding cells were treated with the indicated inhibitors and 24 hours later with citochalasin B for 30 hours. A total of 300-400 cells were analyzed per independent experiment. Statistical analysis of all figures was performed with a two-way ANOVA followed by a Bonferroni post-test (*p<0.05, **p<0.01, ***p<0.001). Data are shown as the average of 2-3 independent experiments with the standard error of the mean.

**Figure 6 - figure supplement 1.**
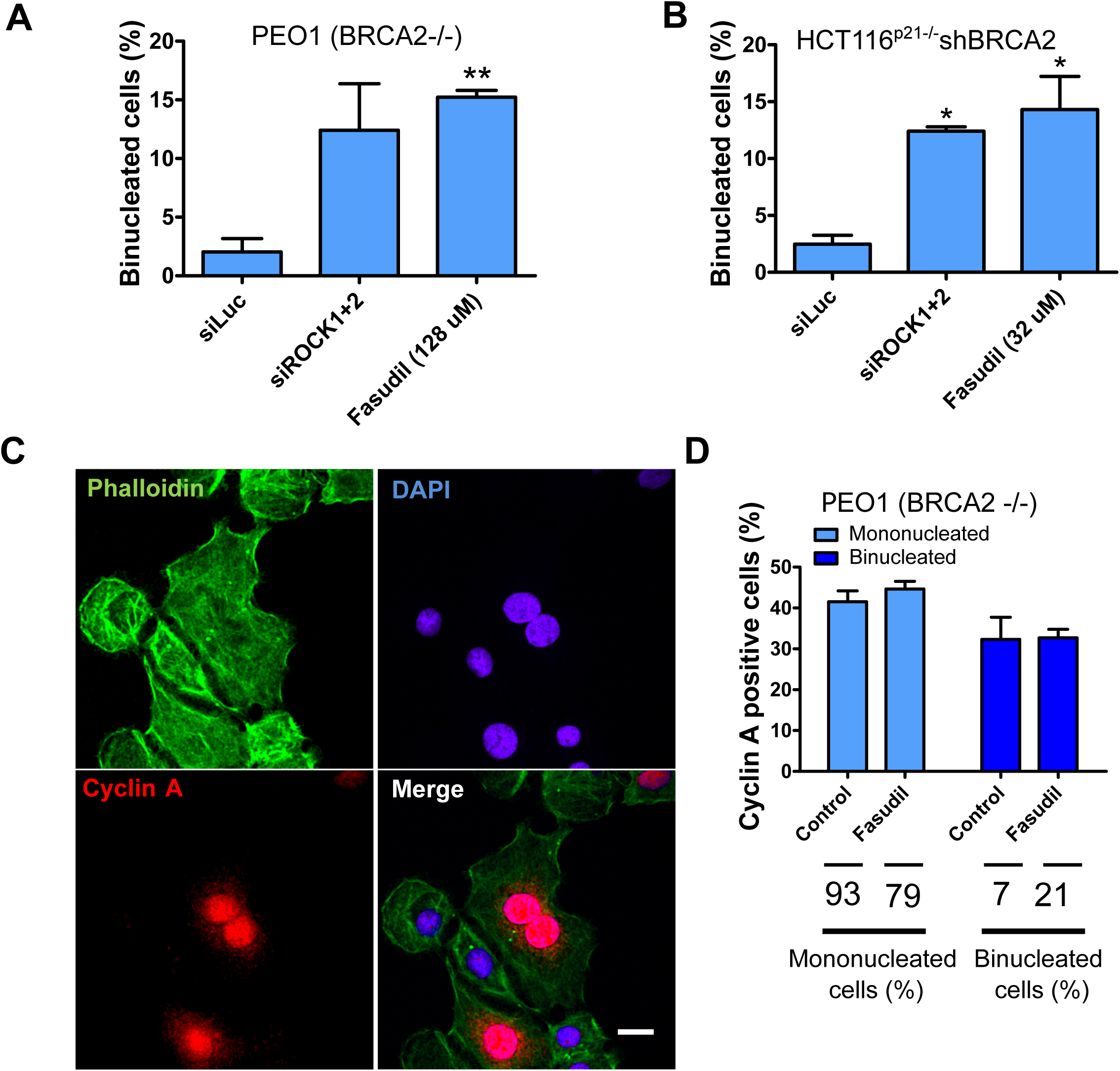
Multinucleated BRCA2-deficient cells resulting from Fasudil treatment are able to transit through S phase. **A)** Percent of binucleated PEO1 cells transfected with siROCK (1+2) or treated with Fasudil (N=2). B) Percent of binucleated shBRCA2 HCT116^p21-/-^ cells transfected with siROCK (1+2) or treated with Fasudil (N=2). **C)** Representative PEO1 cells stained with DAPI, Cyclin A (S phase marker, red) and Phalloidin (actin cytoskeleton, green) after Fasudil treatment (3 days, 128 uM). **D)** Quantification of cyclin A positive cells in each group: mononucleated; binucleated. The % of mononucleated or binucleated /total cells is shown in the lower part of the panel. Statistical analysis was performed using a two-way ANOVA followed by a Bonferroni post-test (*p<0.05, **p<0.01, ***p<0.001). Data in A, B and D are shown as the average of 2 independent experiments with the standard error of the mean.

**Figure 7 - figure supplement 1.**
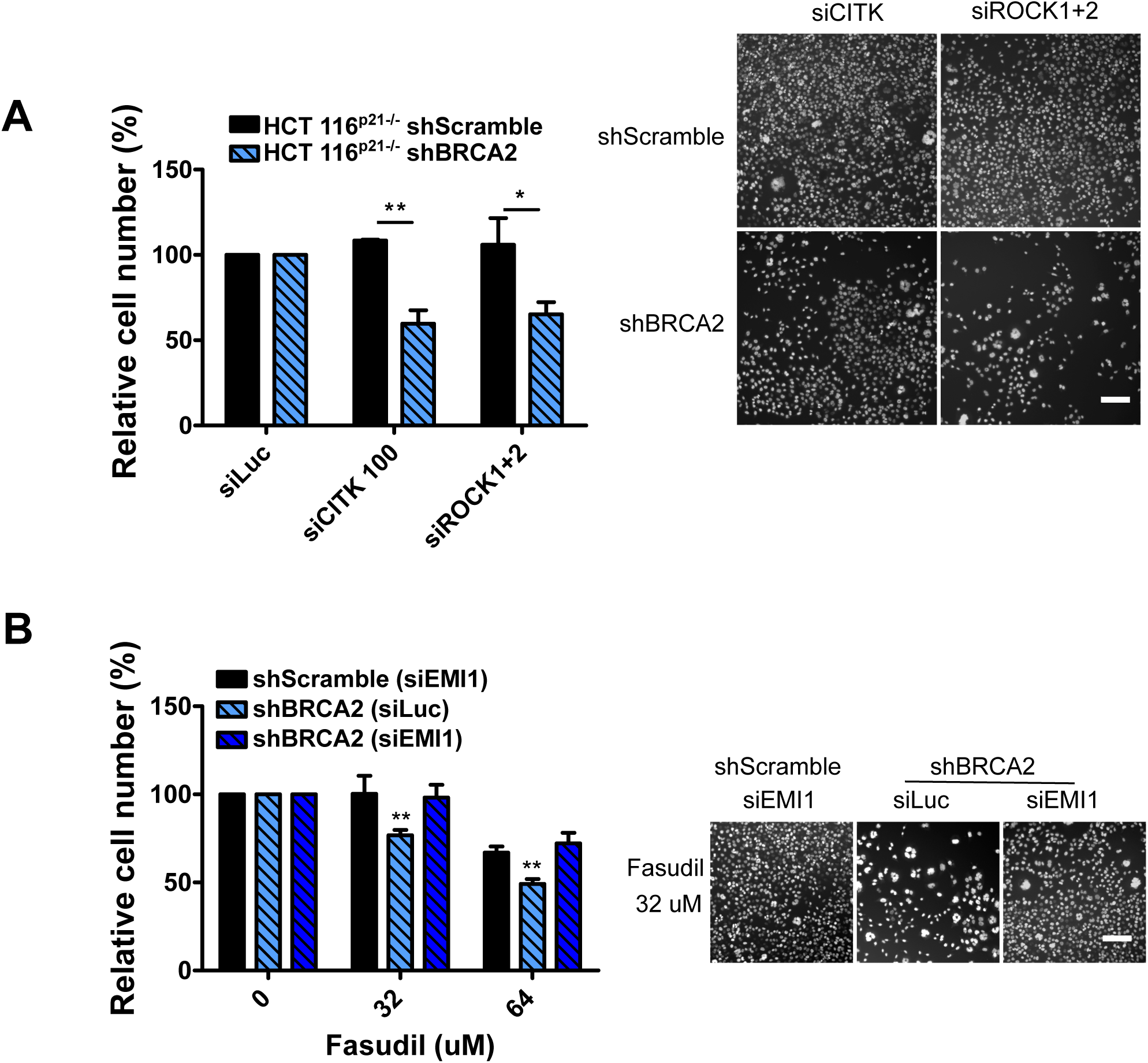
Mitosis as an alternative synthetic lethality strategy for BRCA2 deficient cells. **A)** Relative cell number (%) of shScramble and shBRCA2 HCT116^p21-/-^ cells at 6 days after transfection with siCITK or siROCK (1+2) (N=2). Representative images are shown on the right. B) Relative cell number (%) of shScramble and shBRCA2 HCT116^p21-/-^ cells at 6 days after transfection with siEMI1. Samples were treated with Fasudil when indicated (N=2). Representative images are shown on the right. Statistical analysis was performed using a two-way ANOVA followed by a Bonferroni post-test (*p<0.05, **p<0.01, ***p<0.001). Data are shown as the average of 2 independent experiments with the standard error of the mean

